# Kernel Filter-Based Adaptive Controllers For Cybergenetics Applications

**DOI:** 10.1101/2024.10.21.619394

**Authors:** Benjamin Smart, Lucia Marucci, Ludovic Renson

## Abstract

Cybergenetics is an advancing field that seeks to implement control theory within biological systems. When applying feedback control for the regulation of gene expression or cell proliferation, model-based control strategies can be applied; in this context, online adaptive mathematical models can be used to keep models in tune with the current behaviour of the biological system. Controllers are often constrained by their sampling rate, which is usually relatively low when using microfluidics/microscopy platforms. Current adaptive filters can lead to an inaccurate predictive model when operating with a low sampling rate, leading to sub-optimal control. Here, we propose a kernel filter that can fit model parameters online to produce a more accurate predictive model that can be included within an adaptive model predictive control scheme. The use of the kernel filter is demonstrated in *in silico* and *in vitro* experiments, where we control a synthetic gene oscillator and a P53 oscillator, and observe a synthetic toggle switch. Our results show that the kernel filter outperforms a particle filter when used for parameter estimation in both the predictive model accuracy and when included within an adaptive model-based controller.

## 1 Introduction

Cybergenetics is a recent field coming out of synthetic biology, which applies principles and techniques from control engineering to forward engineer complex phenotypes in living cells, such as gene expression and cell proliferation [1, 2]. Model-based controllers can be used in cybergenetics [3, 4], often offering better performance than model-free strategies.

Parameter estimation is vital for all systems engineering problems that require a mathematical model [5]. Most biological control applications are mainly based on models with fixed parameters, i.e. parameters estimated offline using calibration data sets [6, 7]. However, when conducting closed-loop experiments, the system of interest may be pushed into a new area of the problem space where the identified parameters cannot describe its behaviour [8]. It is well known that mathematical models of biological systems can suffer from identifiability issues [9] due to the impossibility to directly measure most of the parameters and noisy behaviors, in addition to possible uncertainties in the structure of the model [10, 11]. Therefore, it is beneficial that online parameter estimation is included within cellular model-based feedback control schemes.

Online parameter estimation combines the benefits of physics-based and data-driven control, utilising some of the stability guarantees and robustness of the model-based controllers whilst avoiding sensitivity problems due to the uncertainties in fixed model parameters and any noise/variance in observations, [12, 13].

Regarding the experimental implementation, cybergenetics mainly relies on the use of microscopy/microfluidics platforms, where time-lapse imaging is combined with image segmentation [14] to quantify real-time outputs (for example, gene expression), compute the control error by comparing the output to the target behaviour, and finally apply control algorithms which can provide, to cells, control inputs to minimise the control error [15, 16, 17, 18, 19, 20]. The control algorithms can include the online fitting of models, often using particle filters [21]. The sampling rate at which images are collected and analyzed is usually limited due to the physical constraints of the microscope and, most importantly, issues related to phototoxicity when performing long experiments. Therefore, there is a need for adaptive models that can be fitted at a lower sampling rate. Particle filters are frequently used to extract biological system state and parameter information offline, [22, 23] and online [7, 24]. Recently, particle filters are also being used within feedback controllers working on biological systems[25, 21].

Here, we propose a kernel filter that can fit model parameters at low sampling rates and be included within an adaptive model predictive controller (MPC). The kernel filter proposed here is adapted from [26], where parameter estimation was performed offline on a mechanical system. In this paper, we extend this to use the kernel filter online, using a shifting kernel window and a shifting sampling grid to solve the nonlinear optimisation problem online, updating the system parameters and kernel hyperparameters each sampling period.

Many biological systems present nonlinear dynamics such as temporal oscillations and hysteresis, occurring across different temporal and spatial scales [27]. The first example considered here is a synthetic gene oscillator implemented in bacterial cells [28]; we used the mathematical model of the system reported in the original publication to simulate its dynamics, so we will refer to it as an *in silico* oscillator. The second example is the P53 biological oscillator [29]; it is a natural oscillator found in mammalian cells, which controls several critical cellular processes, including cellular senescence, apoptosis, cell cycle arrest, and DNA repair; also, here we used a model of the system to reproduce its dynamics. The third example is a synthetic toggle switch [30], that allows engineered bacteria cells to exhibit two stable steady states of gene expression. In this case, we checked our newly proposed kernel filter for parameter fitting on experimental data. Of note, the bacterial vs mammalian systems we used as test beds present very different dynamics (the bacterial systems are an order of magnitude faster than the mammalian one). We observed that the kernel filter outperforms a particle filter when a low sampling rate (less than forty samples per period for the bacterial systems and three samples per period for the mammalian system) is used. We also show that, for the bacterial systems, the kernel filter is better over all sampling rates.

The structure of this paper is as follows. Section 2 describes the kernel and particle filters, including an overview of the filters used within a feedback controller using MPC. Section 3 compares the performance of the two filters in predicting the future behaviour of the states and compares the performance within an MPC controller for the three genetic networks. Section 4 presents the conclusions of this study.

## 2 Methods

### 2.1 Kernel Filter

Consider the dynamic system

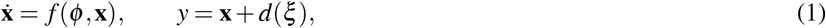

where **x** are the states of the system, *f* is a known nonlinear vector field and *ϕ* are the unknown model parameters. *y* represents the measurement of the states, which is assumed to be perturbed by additive Gaussian noise, *d*, of covariance, *ξ*. The state estimate at time 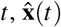, is given by solving

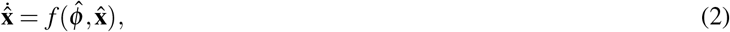

where 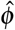 are the estimated model parameters.

Kernels are frequently used to predict a future state [31], by lifting lower dimensional nonlinear problem space measurements, *y*, to infinite dimensional linear feature spaces. The kernel is then used to describe the cross product of two measurements in this Hilbert space, meaning that the infinite transformation dictionary never has to be evaluated. Rather than comparing a single measurement to a model prediction, the kernel can be used to relate all previous measurements and indicate whether the prediction from a candidate model seems reasonable when compared to all the past measurements. In Ref. [26], the authors estimate the nominal parameter set, *ϕ*, by maximising the similarity between a kernel containing the outputs of a candidate model and a kernel containing just the measurements, using the Hilbert Schmidt Independence Criterion (HSIC) [32]. Here, we adapt this method to work online, to be part of an adaptive MPC re-estimating the system parameters within each time step and using a sliding kernel window and an adaptive grid search.

We define the measurement vector, *Y*_*t* −*W*:*t*_ = [*y*^*T*^ (*t* − *W*), *y*^*T*^ (*t* −*W* + 1),…, *y*^*T*^ (*t*)] and a state vector, *X*_*t* − *W*:*t*_ = [**x**^*T*^ (*t* −*W*), **x**^*T*^ (*t* −*W* + 1), …, **x**^*T*^ (*t*)], where *W* represents the kernel’s sliding window size. *W* prevents the kernel from growing in size as more measurements are taken.

The prediction kernel, 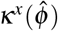, uses a mixed vector, 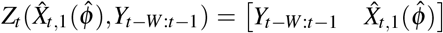 containing past measurements and one predictive step, 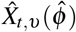, where *υ* is the step number within the prediction horizon used. Following investigations, a Periodic kernel is used in all simulations as discussed in Section S1. Each element of the kernel is calculated as

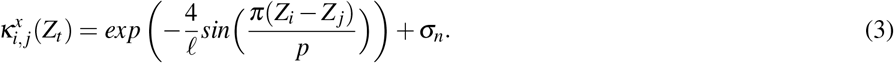

The measurement kernel, *κ*^*y*^, is calculated using only the measurements *Y*_*t*−*W*:*t*_ as

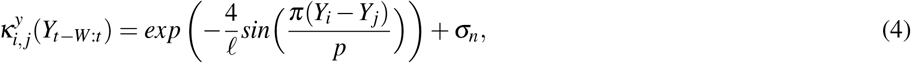

where *p* is the periodic hyperparameter, ℓ dictates the exponential length scale of the kernel and *σ*_*n*_ weighs the expected noise. The hyperparameters are initially chosen by the method described in [26], and then optimised, online, using Marginal Likelihood, [33], throughout the first kernel window. Hyperparameter optimisation is further discussed in Section S2.

The similarity between the predicted kernel and measured kernel spaces is compared using the Hilbert Schmidt Independence Criterion (HSIC) [32],

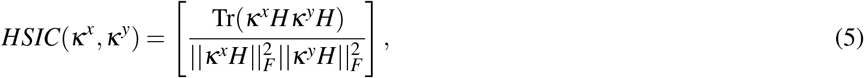

where 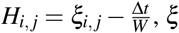 is the covariance of the noise present (assumed to be white noise, *ξ* is a diagonal matrix) and Δ*t* is the time difference between samples. The score tends to one if the two kernels are identical and tends to zero as the difference increases. The global maximum of the HSIC would correspond to the nominal parameter set, *ϕ*.

To find 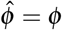, Ref. [26] implements both a grid search and a nonlinear optimiser algorithm. Their grid search is a static offline random sample search. Here, we implement a moving sample window around the operating point, as outlined in Algorithm 1, so the filter can operate online. Gradient-decent-based nonlinear solvers (MATLAB’s® fmincon, IPA, and NOMAD [34]) were not used as it was found that they often settle on local minima near their initiation point, not improving the predictive states.

A grid search requires some knowledge of the initial parameter values to define the finite grid and is otherwise very computationally heavy. The grid used here trials three different values for each parameter, one at the previous optimal parameter value, 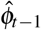, and another two at ±*ρ*%. The possible parameter change between any two samples depends on the sampling rate. Therefore, the search grid intervals, *ρ*, should depend on this sampling rate. This relationship is discussed in Section S3. For a system containing *n* parameters, the discrete set, Φ_*t*_, contains all 3^*n*^ possible combinations of 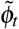 where each element of 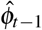 has been multiplied by an element of ([1 −*ρ*, 1, 1 + *ρ*]) to create a 3^*n*^ unique trial 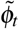. Whichever sample set, 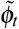, produces the highest HSIC will be chosen as the kernel filter’s parameter estimate, 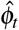. The grid size is defined relative to the parameter values to follow the observed parameter changes. A finer grid can be used, resulting in higher computational costs.

#### Algorithm 1

Kernel Filter

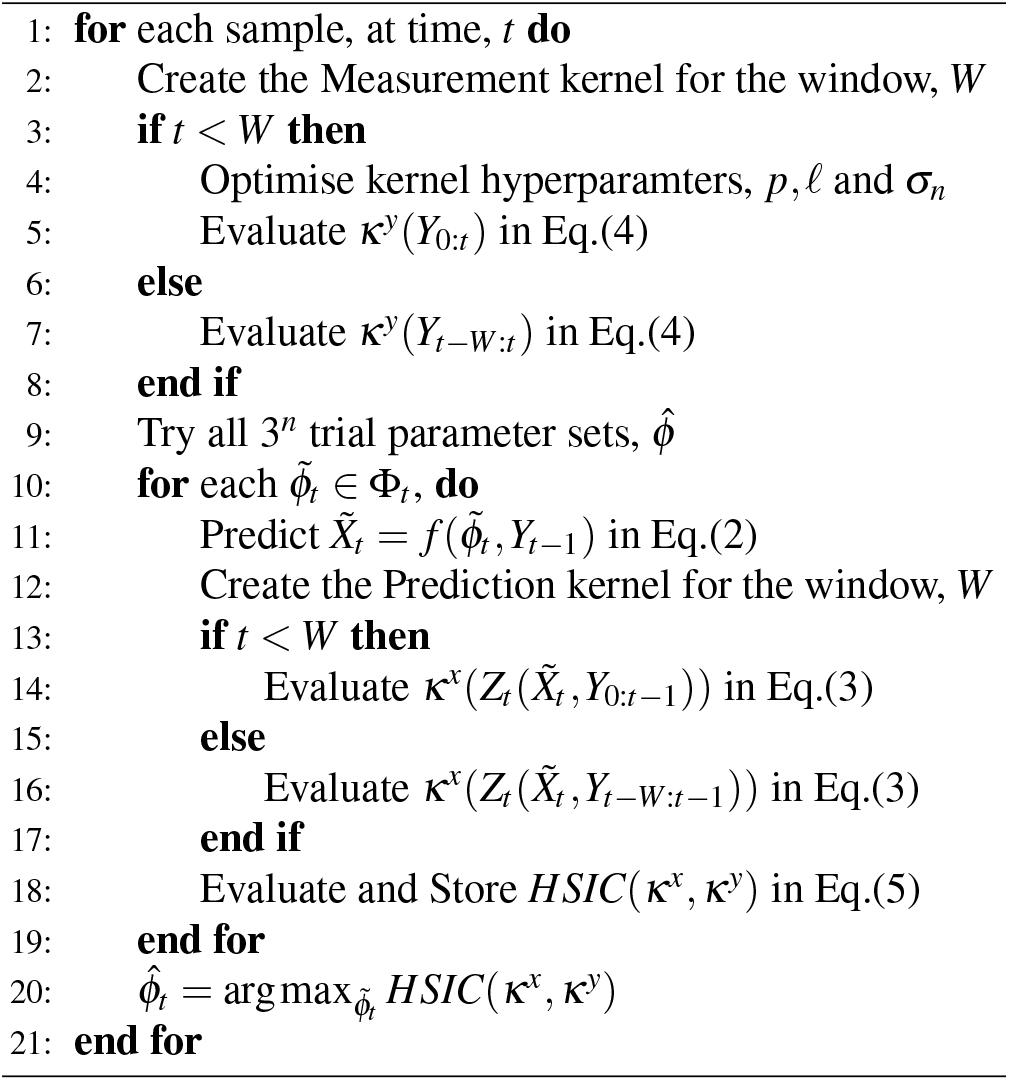

### 2.2 Particle Filter

Particle Filters are a Bayesian filter class that estimates the parameter distributions based on output observations [35]. It is a well-established method for fitting models and is included here for comparison. The particle filter aims to find *P*(*ϕ* |*Y*_0:*t*_), the probability of the parameter set given the past measurements. The first two are the basis of all Bayesian filters, using Bayes’ theorem to state an a priori about the distribution of the parameters, *P*(*ϕ*), and the observations of the output to measure, *P*(*Y*_0:*t*_ | *ϕ*). Marginalisation alongside marginal likelihood is used to approximate the probability of the output, regardless of the parameter set, *P*(*Y*_0:*t*_), by propagating *M* sample parameter sets, 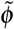, through the system,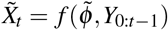. The distribution of the output, *P*(*Y*_0:*t*_|*ϕ*), can be approximated as the distribution of these samples,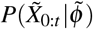. Each parameter set, 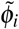, is weighted by *ω*_*i*_, an estimate of how likely its output, 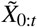 would be to occur for the given output distribution, *P*(*Y*_0:*t*_|*ϕ*). Each *ω* is summed to form an a posteriori parameter distribution,

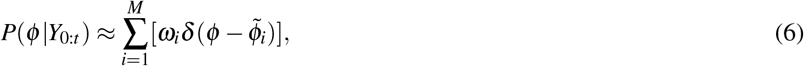

where *δ* is the Dirac distribution. With the estimated parameter set as the expected value of this distribution, 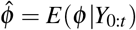.

The final step of a particle filter utilises an importance distribution, scaling the weighting, *ω*, by how accurate our model is perceived to be, knowing that it is not an exact representation of the system. After each iteration, the particles can be resampled. The particle filter simulation parameters are discussed in Section S4.

### 2.3 State Error Index

To assess the error of the predictive model fitted by the filters, we define the identification Mean Absolute Error, *MAE* [36], as

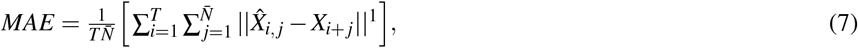

calculating the sum over the total simulated time, *T*, of the normalised absolute difference between the predictive states, 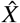, and the actual states *X*, over an *N* step prediction horizon. 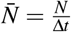, the total number of samples within the N-step horizon.

### 2.4 Feedback Control Method - Adaptive Model Predictive Control (MPC)

MPC uses a model of the plant (system to be controlled) in a feedback loop to estimate the effect of future control inputs on the states, *N* time steps into the future. Prediction horizon steps, *N*, are discrete time steps in the unit of the model (minutes for the bacterial systems in Sections 3.1 and 3.3 and hours for the mammalian system in Section 3.2), evaluating the predicted state at each step. These control inputs are chosen to minimise a cost function, *J*(**E, U**), along with any control input or state constraints [37],

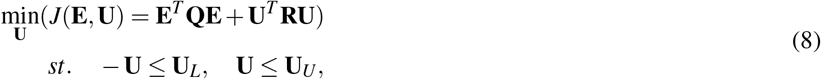

MATLAB’s® nonlinear solver, ‘fmincon’, solves the nonlinear optimisation problem with its interior point algorithm, limited to ten iterations. **E** = [**e**(0), **e**(1),…, **e**(*N*)]^*T*^ describes the future predicted state errors using control inputs, **U** = [**u**(1), **u**(2),…, **u**(*N*)]^*T*^ in the estimate model of the system. The updated model parameters are provided by either the kernel filter or the particle filter in each iteration (outlined in Section 1). Therefore, the model within the MPC is fitted online to the observations being made.

The weight of each cost function term can vary the optimal control input. In the examples described here, the control inputs have to stay within their bounds, and only the tracking error is costed (**R** = 0, and **Q** = 1). In all examples, up to the first ten future control inputs are applied to the system to reduce the computation cost and increase the effect of the model on control performance.

**Figure 1.**
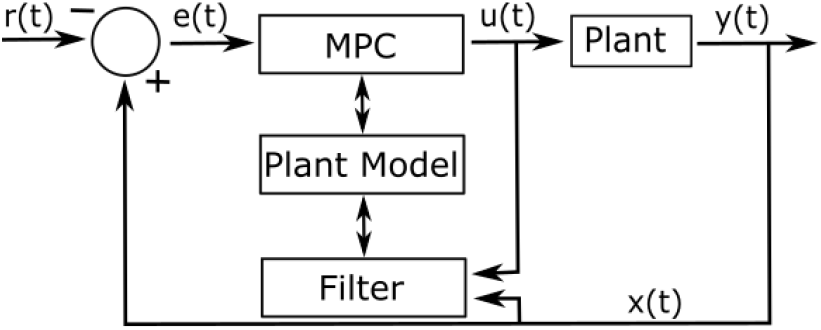
The MPC architecture. The biological system to be controlled, the plant, is observed in the output, *y*. The model states, *x*, are assumed to coincide with the plant outputs *y. x* is given to the filter to fit the plant model and compared to the reference, *r*, to be given to the MPC algorithm. The MPC optimally chooses the input, *u*, which acts on the plant system.

### 2.5 Control Error Index

The performance of the controller is taken as the error between the output state and its reference; we define the control Root Mean Squared Error, *RMSE*, as

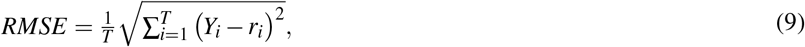

calculating the sum over all time, *T*, of the difference between the actual output *Y* and reference, *r*_*i*_ = *r*(*t* = *i*Δ*t*).

## 3 Results

### 3.1 Synthetic Gene Oscillator

The model of synthetic gene oscillator found in [28] has been used here, describing the interactions between a CI protein (*X*_1_) and a LAC protein (*X*_2_). A four-state kinetics model, containing both the proteins and their corresponding dimers, is reduced to a two-state model containing a dimensionless representation of the CI protein (*x*_1_) and the LAC protein as (*x*_2_),

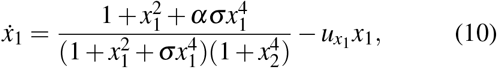

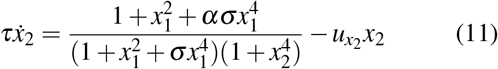

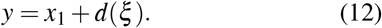

The CI protein promotes both the CI and LAC dimers, whereas the LAC protein inhibits both dimers, as shown in Fig. 2. The system parameters are discussed in Section S5. For all time varying parameter simulations, *ϕ*_0_ = [*α, σ*] = [11, 2].

**Figure 2.**
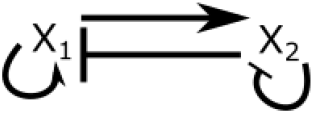
The synthetic gene oscillator in [28]. The schematic shows the interactions between a CI protein (*x*_1_) and a LAC protein (*x*_2_); arrow-ended lines indicate activation, and bar-ended lines indicate inhibition.

#### 3.1.1 Low sampling rate filtering

To compare the error of the kernel filter to the commonly used particle filter, [38], *α* and *σ* within the nominal parameter set, *ϕ*, have been changed throughout a simulation. This change can be seen by the black lines in Fig. 3B, with an increase in *α* to 2.25 times its initial value, then an equivalent decrease, followed by an increase of *σ* to 2.25 times its initial value, then an equivalent decrease. Within the nominal parameter set, *ϕ*, all other parameters remain at their initial values throughout the simulation. The filters attempt to fit the parameter set, 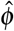, such that state predictions, 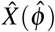, follow the actual states, *X* (*ϕ*). The prediction trajectories show the state trajectories that are predicted using the state and estimated parameters. Predictions are made at each output sample.

**Figure 3.**
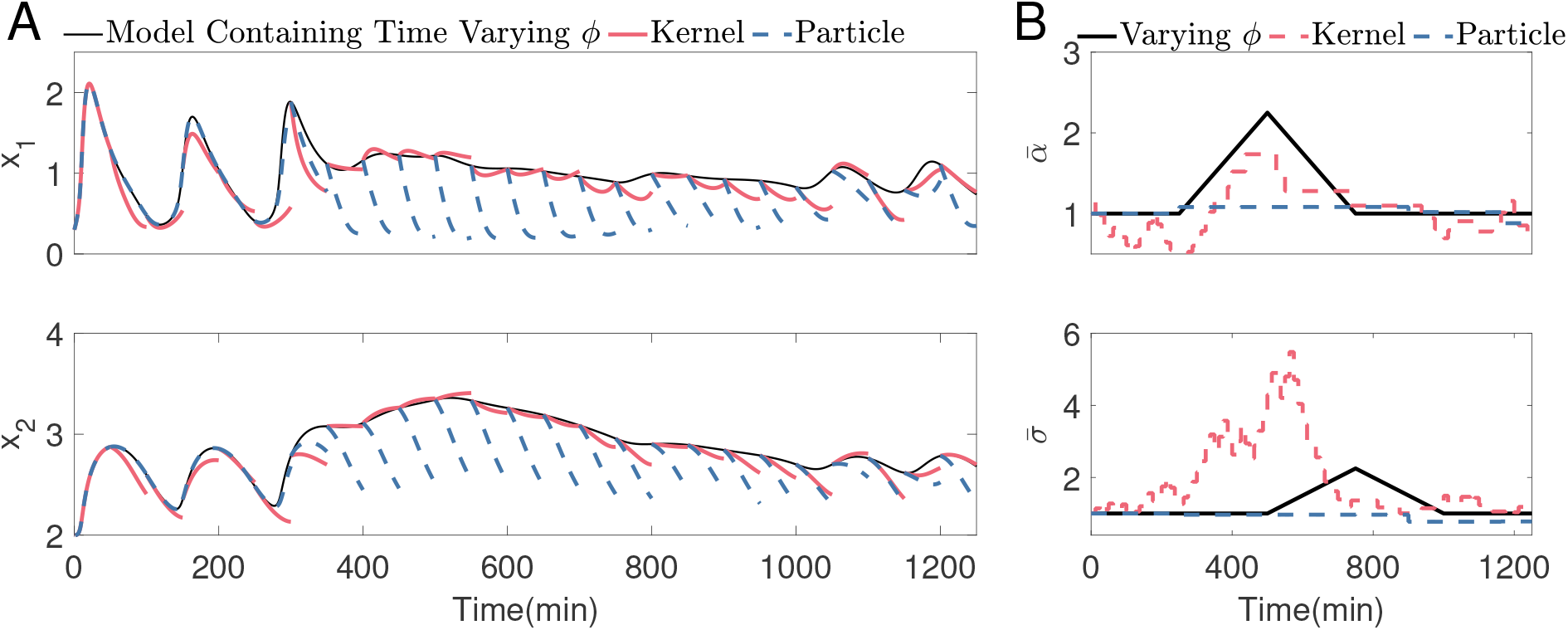
The filters fitting the parameters, online, to match states of the synthetic gene oscillator (Eq.(10) - Eq.(11)) in which the nominal parameter value, *ϕ*, changes over time. One sample every 12.5 minutes (8.8 samples per period) is used, which is considered a low sampling regime here. Other simulation parameters: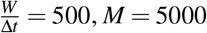, 0.5 re-sampling threshold, 0.0158 expected variance of parameters. A) Every 50 minutes, the prediction trajectory, 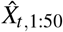 of each filter is compared to the nominal model. B) The parameter fitting of 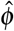, compared to the nominal parameters, *ϕ*. Each parameter has been normalised by the initial parameter defined in Section 3.1.

Fig. 3A displays the predicted trajectories when the system is sampled every 12.5 minutes. The kernel filter manages to change the amplitude and period of the oscillations between 300 and 900 minutes to almost no oscillations. In contrast, the particle filter struggles to remain close to the system’s response simulated with nominal parameter values. The identification *MAE* reflects this as 0.0884 *<* 0.2850 for the kernel and particle filter, respectively.

Fig. 3B shows the parameter variation. It can be seen that the kernel filter makes up for the decrease in *α* by initially increasing *σ*, still achieving a reasonable state estimation. It is also shown that the particle filter does not significantly change either parameter from the initial estimation. The model is over-parameterised, resulting in parameter non-identifiability [39], meaning that there are several parameter sets, 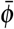, that lead to near identical state predictions and therefore, the filter could choose any of these sets as 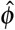 and still achieve good state tracking. Section S6 includes simulations with measurement noise of covariance *ξ* = 10^−2.5^.

#### 3.1.2 Low sampling rate control

As discussed in Section 2.4, either filter can be used to fit the model within an MPC scheme. The synthetic gene oscillator has been synthetically designed to oscillate in its free response [40]; the controller discussed here can be used to steer the amplitude and period of the oscillations. A sine wave with a frequency of roughly ten times slower than the free response is used as the reference, **r**(*t*). These slow reference dynamics can highlight any errors within the MPC’s model.

Identical to Section 3.1.1, the temporal evolution of *α* and *σ* within the nominal parameter set, *ϕ*, has been changed throughout a simulation, shown by the black lines in Fig. 4B. The filters attempt to fit the parameters, 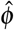, such that state predictions, 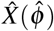, follow the actual states, *X* (*ϕ*) and therefore give the MPC algorithm a more accurate model.

**Figure 4.**
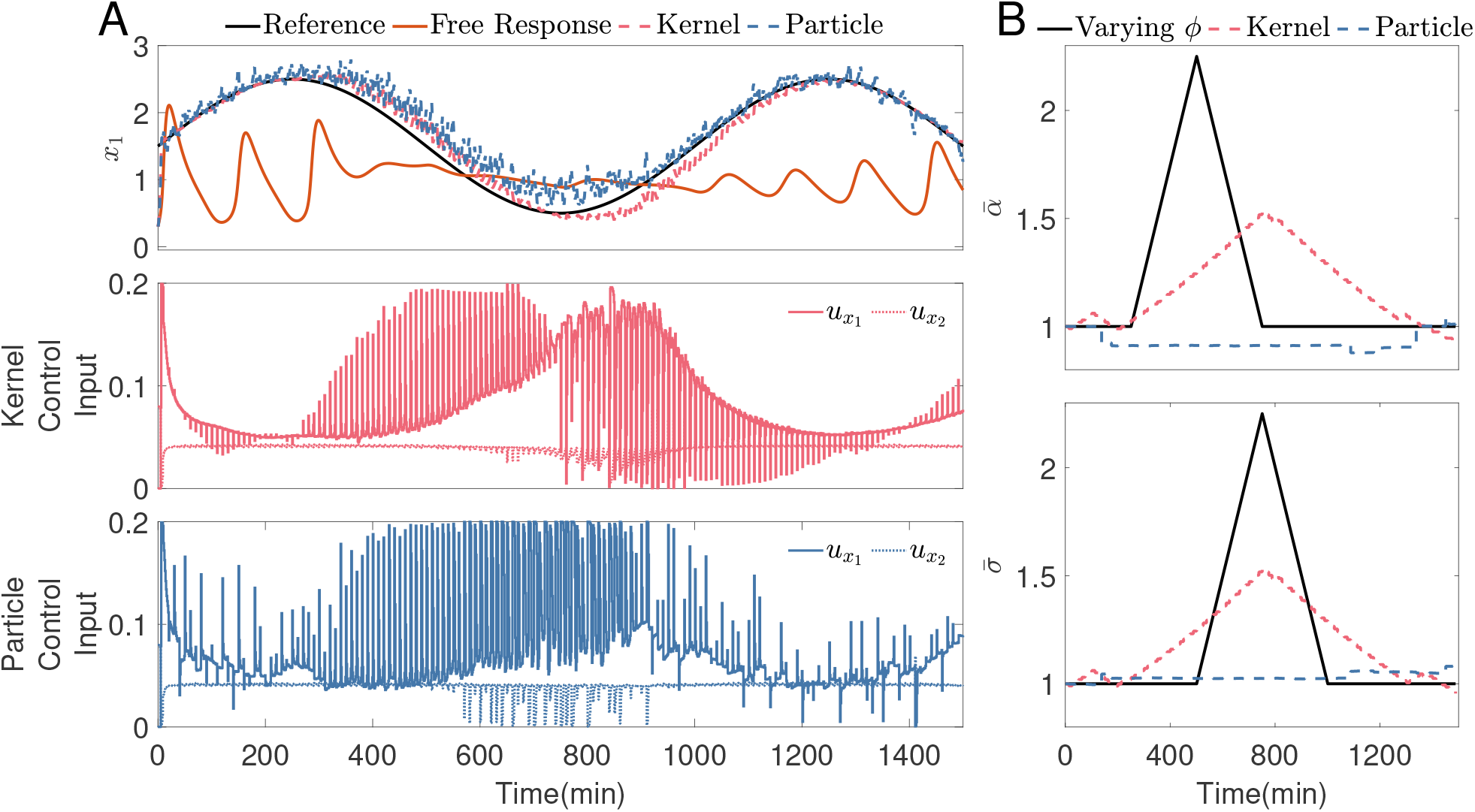
The filters fitting the parameters, online, to match the states of the synthetic gene oscillator (Eq.(10) - Eq.(11)) in which the nominal parameter value, *ϕ*, changes over time. One sample every 12.5 minutes (8.8 samples per period) is used, which is considered a low sampling regime here. Other simulation parameters: the control input is actuated once a minute; *N* = 10 minutes, containing ten input steps 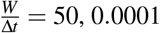 is the expected variance of parameters, *M* = 5000, 0.5 resampling threshold. A) The system’s output qualitatively follows the sine wave reference using the control input shown. B) The parameter fitting of 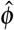, compared to the nominal parameters, *ϕ*. Each parameter has been normalised by the initial parameter defined in Section 3.1.

Fig. 4A shows the synthetic gene oscillator control response sampling every 12.5 minutes (8.8 samples per period). It can be seen that when the free response amplitude and centre of oscillation change between 400 and 1000 minutes, the kernel filter has a smaller control error than the particle filter. The particle filter’s MPC believes the system would naturally oscillate with larger amplitudes between these times, causing the controller to miss the reference. The control *RMSE* of the synthetic gene oscillator is 0.1529 for the kernel filter and 0.2330 for the particle filter.

Fig. 4B shows the parameter variation of the synthetic gene oscillator sampling every 12.5 minutes. The particle filter rarely changes the parameters, whereas the kernel filter increases *α* and *σ* when *α* increases in the nominal set. These parameter changes lead to the smaller control error seen in Fig. 4A.

#### 3.1.3 Varied sampling rate identification and control

The identification Mean Absolute Error *MAE* Eq.(7) of the two filters is compared over a range of sampling rates in Fig. 5A, looking at the prediction trajectories (as in Fig. 3). The control Root Mean Squared Error, *RMSE* Eq.(9) of the two filters is compared over a range of sampling rates in Fig. 5B comparing the control performance of the two filters following the same reference as described in Section 3.1.2, used within MPC controllers (as in Fig. 4). It can be seen that the kernel filter outperforms the standard particle filter over all sampling rates up until 80 minutes (1.57 samples per period), where both filters follow the identification *MAE* of the static model.

**Figure 5.**
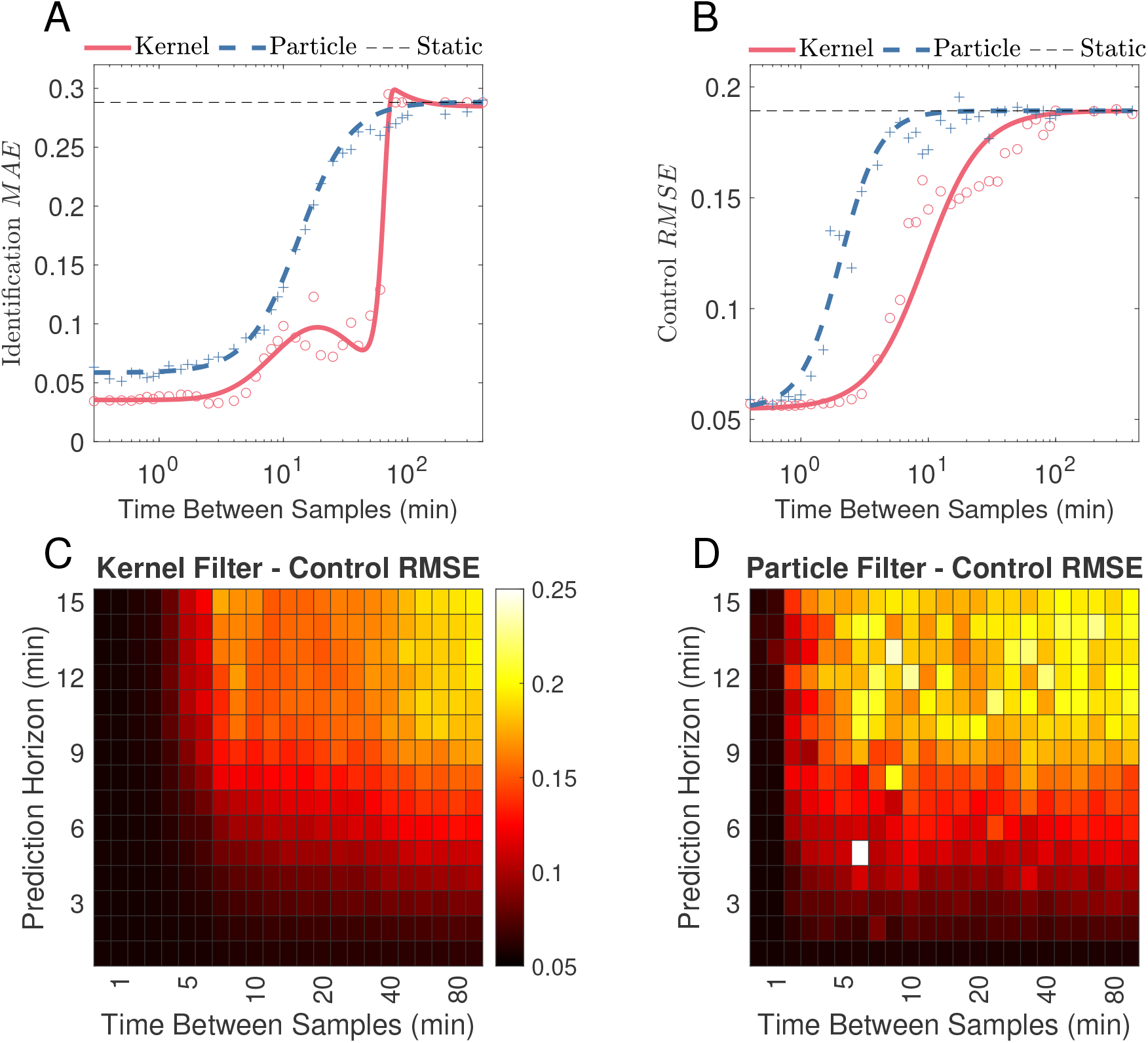
A) The identification Mean Absolute Error (*MAE*, Eq.(7)) of the two filters is compared over a range of sampling rates for the synthetic gene oscillator. Fig. 3 notes the simulation parameters for the kernel and particle filters. B) The control Root Mean Squared Error (*RMSE*, Eq.(9)) of the two filters is compared over a range of sampling rates for the synthetic gene oscillator, with simulation parameters as in Fig. 4. C) and D) The control *RMSE* of the kernel filter(C) and the particle filter(D), compared over a range of sampling rates and prediction horizons, *N*, for the synthetic gene oscillator. Other simulation parameters are in Fig. 4.

Similarly to the identification MAE in Fig. 5A, the control *RMSE* in Fig. 5B shows the kernel filter outperforming the particle filter over all sampling rates.

For fast sampling rates, it is expected that the identification *MAE* and control *RMSE* would be reduced as the filters are observing the system more frequently. This can be seen for both the particle and the kernel filter. Once the filter is used in a rapid sampling regime, the best filter will depend on the filter setup and the underlying system itself. Depending on the relative size of the parameters and the output’s sensitivity to each parameter.

The control *RMSE* can be seen in Fig. 5C and D over varied sampling rates and prediction horizons, *N*. For both the particle and kernel filters, it can be seen that as the prediction horizon gets longer, the control error increases. We expect that there will be a decrease in tracking error with longer prediction horizons as the controller gains more information about the future consequences of the inputs. However, for nonlinear systems with time-varying parameters, longer predictions with fixed parameters, 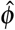, can mislead the MPC [41]. Model prediction trajectories consider fixed parameters, 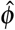, which are only locally accurate. A significant decrease in control *RMSE* can be seen when including a second time step in the prediction horizon for slower sampling times. This is explained by the fact that each prediction step represents a significant jump in time for each state at such a low sampling rate. For instance, for 80 minutes between samples, one prediction step represents almost 0.75 times the oscillation period of the system.

As the sampling rate decreases, the control error increases. Therefore, we expect the bottom left corner of the plot to be the darkest region, with the lowest control error and the top right corner to be the brightest, with the highest control error. For the kernel filter in Fig. 5C, for all prediction horizons, the offset of the darker region persists until larger sampling rates are compared to the particle filter in Fig. 5D. There is a significantly darker region for the kernel filter compared to the particle filter between sampling times of 2.5-60 minutes (44 − 1.83 samples per period) and any prediction horizon, showing that the kernel filter outperforms the particle filter, not only at *N* = 10 (as in Fig. 5B) but for all prediction horizons shown.

### 3.2 P53 Oscillator

The model of a P53 oscillator describes the interactions between the p53 protein and the Mdm2 protein. Model 4 from [29] is used here, containing three states: P53 protein as *x*_1_; the Mdm2 precursor as *x*_2_ and the Mdm2 protein as *x*_3_.

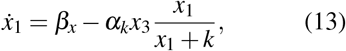

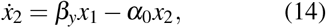

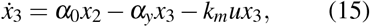

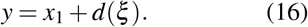

A schematic of the P53-Mdm2 interaction is shown in Fig. 6, displaying this cascade system’s feedback through an inhibition of P53 from Mdm2. The input to the system, *u*, is an extra degradation of Mdm2 [42].

**Figure 6.**
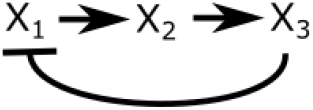
P53 Oscillator formed by negative feedback from an Mdm2 protein (*x*_3_) on a P53 protein (*x*_1_), including a transcriptional Mdm2 precursor (*x*_2_). Arrow-ended lines indicate activation, and bar-ended lines indicate inhibition.

The system parameters are discussed in Section S7. For all time varying parameter simulations, *ϕ*_0_ = [*α*_*y*_, *α*_0_, *α*_*k*_, *k, β*_*x*_, *β*_*y*_] = [0.8, 0.9, 1.7, 0.001, 0.9, 1.2]. The system has six kinematic constants, and three have been selected to be fitted within the filters. *α*_0_, *α*_*k*_ and *β*_*y*_ have been chosen as they characterise the speed of the two cascade reactions, *x*_1_ → *x*_2_ and *x*_2_ → *x*_3_ as well as the feedback inhibition, *x*_3 ⊣_ *x*_1_.

#### 3.2.1 Low sampling rate filtering

We compare the state trajectory error of the filters as discussed in Section 3.1.1. This change in *α*_0_ and *α*_*k*_ can be seen by the black lines in Fig. 7B, with the increase in *α*_0_ to 1.5 times its initial value then an equivalent decrease, followed by a decrease of *α*_*k*_ to 0.5 times its initial value, then an equivalent increase. The prediction trajectories are the future predictions using the current model, taken from the point where every fourth measurement is taken. The trajectories are 10 hours long.

**Figure 7.**
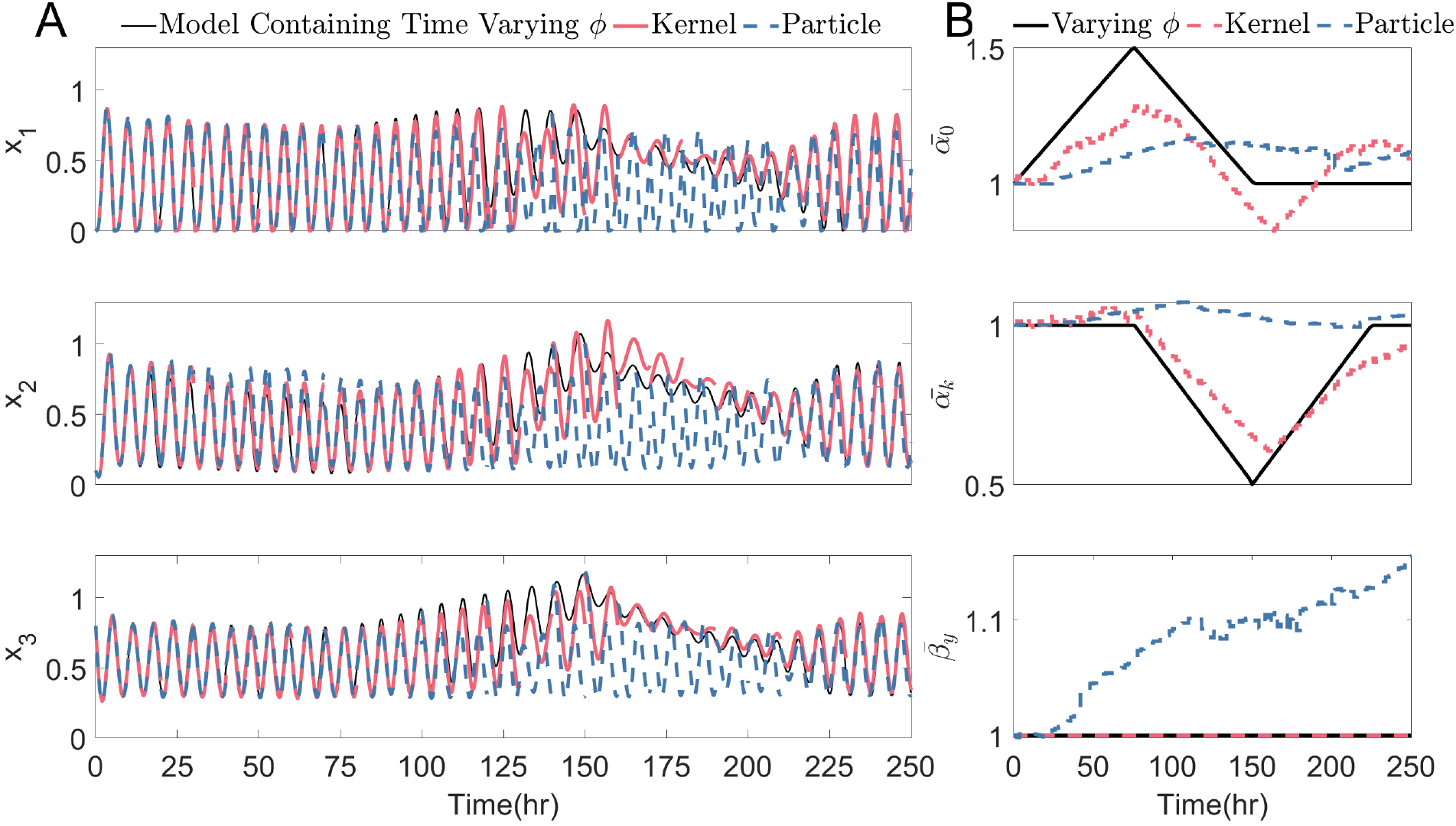
The filters fitting the parameters, online, to match the states of the P53 system (Eq.(13) - Eq.(15)) in which the nominal parameter value, *ϕ*, changes over time. A sample every 2 hours (3 samples per period) is used, which is considered a low sampling regime here. Other simulation parameters: 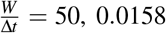 is the expected variance of parameters, *M* = 5000, 0.5 re-sampling threshold. A) Every 10 hours, the prediction trajectory, 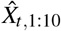 of each filter, is compared to the nominal model. B) The parameter fitting of 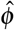, compared to the nominal parameters, *ϕ*. Each parameter is normalised by the initial parameter defined in Section 3.2.

The predicted states of the P53 system, sampled every 2 hours (3 samples per period) in Fig. 7A, demonstrate that the kernel filter outperforms the particle filter. The trajectory length is 10 hours (every fifth measurement), to increase the filter error and make a clear, visible distinction between the two filters’ performances. This can be seen between 25 and 100 hours with the decrease and then increase in amplitude of *x*_2_ and between 125 and 225 hours, where neither filter perfectly matches the changes in the centre of the oscillation for all three states, but the kernel filter does qualitatively and quantitatively better. It can later be seen that this difference is significant in the control simulations between these times. The identification *MAE* (Eq.(7)) of the kernel filter, 0.0685, is significantly less than the identification *MAE* of the particle filter, 0.1760. Section S8 includes simulated measurement noise.

The corresponding parameter variations are shown in Fig. 7B. It can be seen that the kernel filter has minimal variation in *β*_*y*_ whilst following the overall trend of *α*_0_ and *α*_*k*_. The particle filter fails to track these parameter changes, veering all three parameters away from the initial estimate without following the trend of the nominal varying parameters, *ϕ*.

#### 3.2.2 Low sampling rate control

The P53 system is said to synchronise the cellular clock [43], controlling the proliferation cycle. Even in the presence of genetic mutations, this controller could control the amplitude and period of these oscillations to steer cellular senescence, apoptosis, cell cycle arrest and DNA repair [44], leading to effective personalised cancer treatments and regulation of cellular stress. A sine wave with a frequency of roughly ten times slower than the system’s free response is used again as the reference, **r**(*t*). The changes in *α*_0_ and *α*_*k*_ are identical to those discussed in Section 3.2.1.

The control performance of both filters can be seen in Fig. 8A, sampling every 2 hours (3 samples per period). Between 125 and 225 hours, the parameter changes cause the free response to change its centre of oscillation and amplitude. Both filters fit their models to keep up with this, but it can be seen that the particle filter does not keep up with these changes, causing a large error from the reference between these times. The kernel’s control *RMSE* of 0.2297 is less than the particle filter’s control *RMSE* of 0.3255.

**Figure 8.**
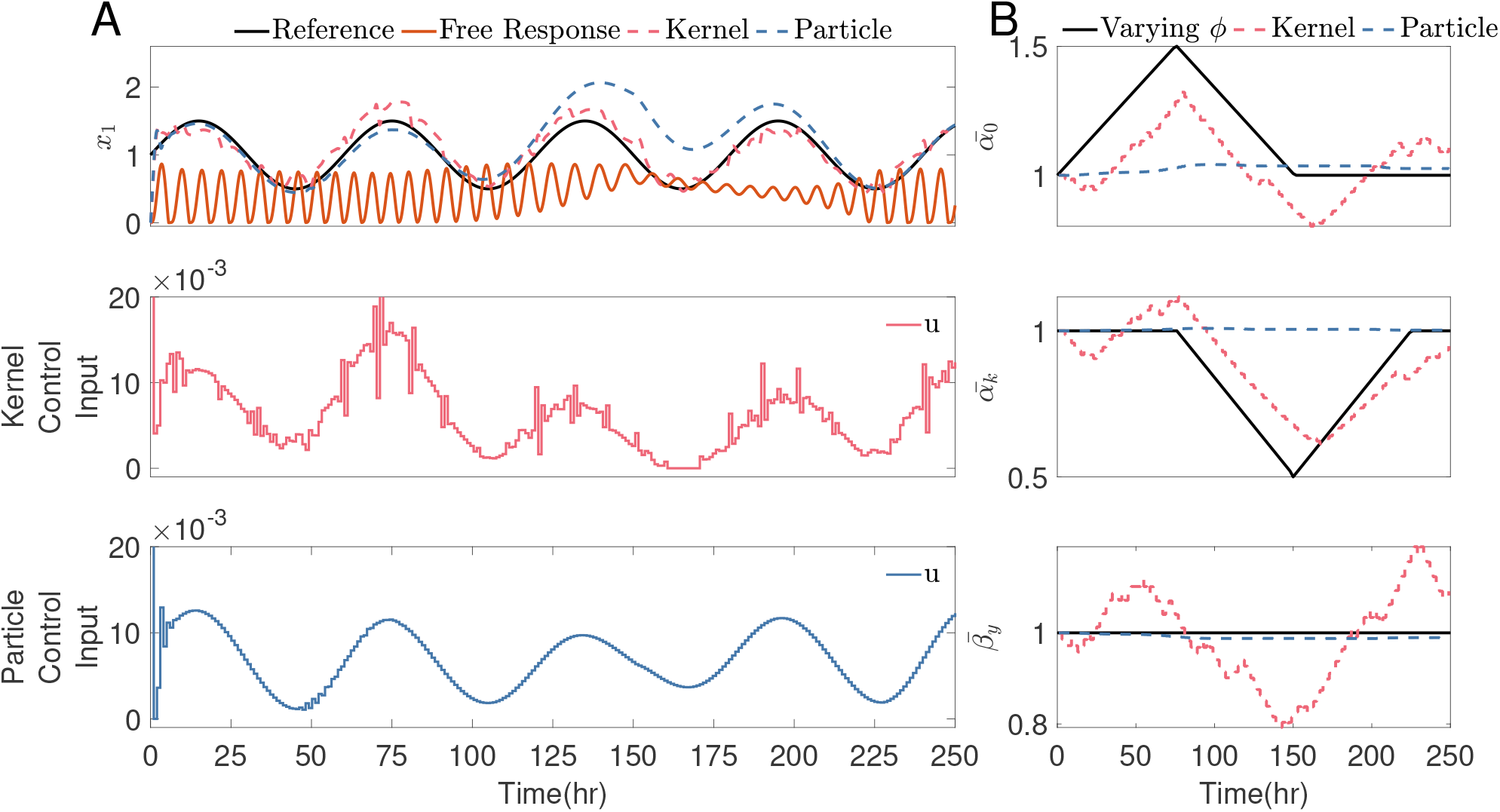
The filters fitting the parameters, online, to match the states of the P53 system (Eq.(13) - Eq.(15)) in which the nominal parameter value, *ϕ*, changes over time. A sample every 2 hours (3 samples per period) is used, which is considered a low sampling regime here. Other simulation parameters: the control input is actuated once an hour; *N* = 10 hours, containing ten input steps; 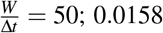 is the expected variance of parameters; *M* = 5000, 0.5 resampling threshold. A) The system’s output qualitatively follows the sine wave reference using the control input shown. B) The parameter fitting of 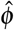, compared to the nominal parameters, *ϕ*. Each parameter has been normalised by the initial parameter defined in Section 3.2.

The parameter changes within Fig. 8B for the P53 system show the kernel filter roughly follows the nominal parameter changes, *ϕ*. Initiating changes in *α*_0_ and *α*_*k*_ to match the time-varying *ϕ*. The particle filter does not have an informed variation from the initial estimation at this sampling rate and remains close to the initial estimation throughout the simulation.

#### 3.2.3 Varied sampling rate identification and control

The identification Mean Absolute Error, *MAE* Eq.(7), indexing the error in the prediction trajectories (as in Fig. 7) and the control Root Mean Squared Error, *RMSE* Eq.(9), indexing the controller performance (as in Fig. 8, using the reference described in Section 3.2.2) are compared over a range of sampling rates in Fig. 9.

**Figure 9.**
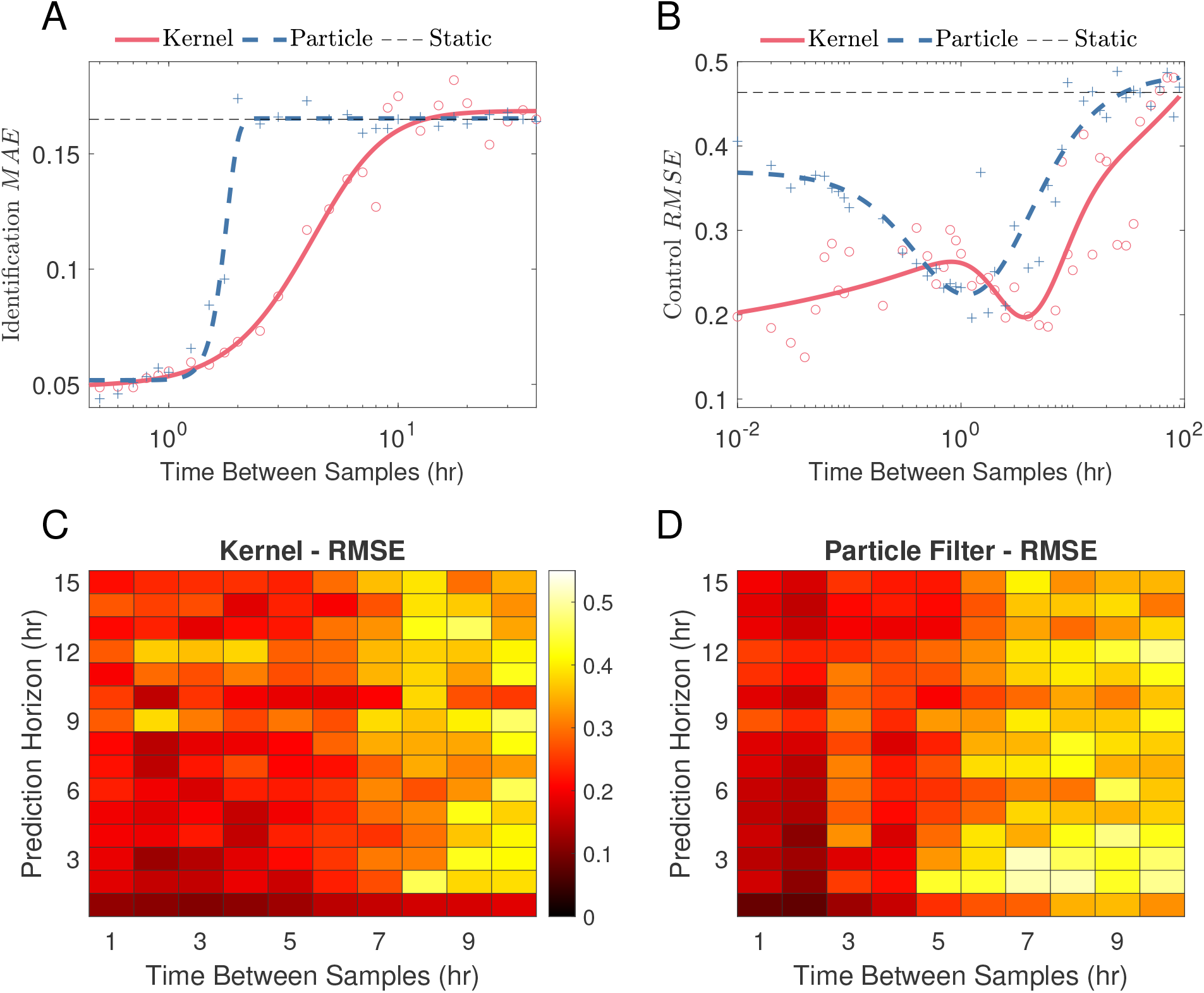
A) The identification Mean Absolute Error (*MAE*, Eq.(7)) of the two filters is compared over a range of sampling rates for the P53 oscillator. The simulation parameters for the kernel and particle filters are noted in Fig. 7. B) The control Root Mean Squared Error (*RMSE*, Eq.(9)) of the two filters are compared over a range of sampling rates for the P53 oscillator, with simulation parameters as in Fig. 8. C) and D) The control *RMSE* of the kernel filter (C) and particle filter (D), compared over a range of sampling rates and prediction horizons, *N*, for the P53 oscillator. Other simulation parameters as in Fig. 8.

For the state trajectories in Fig. 9A, there is a separation of the two filter’s identification *MAE* at a sample time of 1.25 hours (5.14 samples per period), where the kernel filter tracking error remains significantly below that of the particle filter, showing that when sampling less than every 1.25 hours, the kernel filter outperforms the particle filter. Sampling less than once every 10 hours (0.6 samples per period), both filters tend to the identification *MAE* of a static model where the parameters are fixed at the initial value. At this point, neither filter varies the parameters, and the initial steady-state oscillation is unaltered by either filter.

There is a transition at 2 hours between samples (3 samples per period), where the control *RMSE* of the kernel filter crosses that of the particle filter for the first time, showing that the kernel filter outperforms the particle filter for sampling times slower than every 2 hours. This transition point (sampling 1.25 times an hour) is similar to the filtering simulation shown in Fig. 9A.

Fig. 9C and D display the control *RMSE* over varied sampling rates and prediction horizons, *N*. We expect the bottom left corner of the plot to be the darkest region with the lowest control error. There is a darker region for the kernel filter compared to the particle filter between 4 and 7 hour sampling times (1.5-1.17 samples per period) over all prediction horizons, but not between 1 and 3 hours. It can be seen in Fig. 9B that the kernel filter still performs better at slower sampling times, not shown in Fig. 9C and D.

The range of sampling times has been chosen to reflect the likely sampling times in-vitro, avoiding phototoxicity of faster sampling times and the slowest sampling times that are well within the device’s capabilities. In-silico, we find that the largest performance advantages of the kernel filter are at longer sampling times, up to 40 hours between samples, as discussed in Section S9. This demonstrates the theoretical advantages of the kernel filter within adaptive MPC schemes at even longer sampling times.

Fig. 9C is not as smooth as the synthetic gene oscillator in Fig. 5C; multiple ‘light spots’ for both the kernel and particle filters can be seen. This difference is due to P53 system dynamics and the increased number of fitted parameters, as discussed in Section S10. The ODE system has six kinematic constants, yet only three have been fitted here. The more parameters added, the lower the average performance error, but the larger the control error variance, as discussed in Section S11. The grid search limits the parameter space that the kernel filter can jump between each step. Therefore, the kernel filter can be trapped at a local minimum, which causes performance to degrade progressively. Without a more global optimisation algorithm, the filter cannot leave this neighbourhood to retrieve more optimal solutions, leading to the ‘light spots’. A more extensive grid can reduce a ‘light spot’ found in Fig. 9C at *N* = 9, sampling time of 2 hours, from *RMSE* = 0.3473 to *RMSE* = 0.2365 (discussed in Section S12).

### 3.3 Toggle Switch

The toggle switch is one of the most well-studied networks in synthetic biology [45, 46]. We will focus here on the recent implementation in *E. coli* cells proposed in [45], where the network is composed of two proteins, TetR and LacI, that inhibit each other. Two external inputs (i.e. drugs IPTG and aTc) can be used to control the system and make it switch from one stable steady state to the other (Fig. 10). The differential equation model of the system [45] is reported below Eq. (17) to Eq. (22). The first four equations in the model describe the dynamics of TetR and LacI mRNAs and proteins, using mass action kinetics. The final two equations describe the dynamics of the two inputs, differentiating between their extracellular and intracellular concentrations. The system parameters are discussed in Section S13.

**Figure 10.**
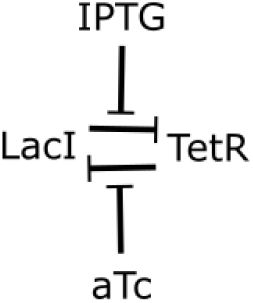
Toggle switch formed by a feedback loop containing two inhibitions between the proteins, LacI and TetR. Each inhibition is inhibited by an extracellular input actively transported within the cell.

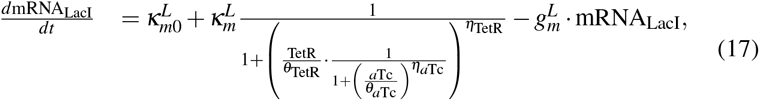

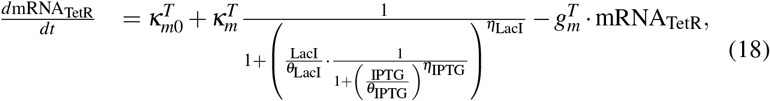

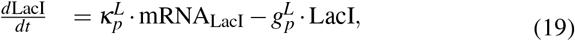

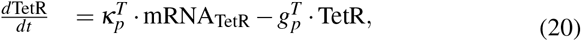

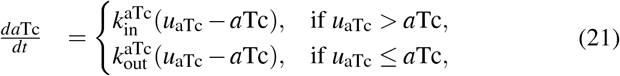

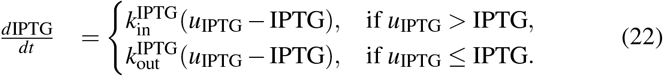

Of the 22 kinematic parameters within the toggle switch system, four have been selected as fitted parameters, *ϕ* = [*η*_*Lac*_, *η*_*Tet*_, *θ*_*Lac*_, *θ*_*Tet*_], as discussed in Section S13. These four parameters are the Hill’s parameters of the two mRNA states, describing the effect of the protein-to-protein inhibition. For all time varying parameter simulations, *ϕ*_0_ = [*η*_*Lac*_, *η*_*Tet*_, *θ*_*Lac*_, *θ*_*Tet*_]= [2.00, 2.00, 31.94, 30.0].

#### 3.3.1 Low sampling rate filtering

The above results used simulated, *in silico* data to compare the performance of the two filters for model fitting, and then used within MPC. Here, we aimed at testing our method on experimental data: we compared the state predictions of the two filters to experimental *in vitro* data from a toggle switch that has been forced to oscillate.

The data is taken from [30], where fourteen experiments, referred to as Exp1 to Exp14, were conducted. The data used can be seen in Section S14. As described in [30], *E. coli* cells embedding the toggle switch gene network were grown in a microfluidic device, allowing for single cell measurement and dynamic provision of media with/without the inducers IPTG and TetR.

We now have an *in vitro* plant system with parameters that change over time. Contrary to the previous *in silico* results, in Sections 3.1.1 and 3.2.1, the controller model is now imperfect/uncertain and the data is corrupted by noise.

In Fig. 11, we compare the state trajectory error of the filters to that of the average cell response within Exp5 (Fig. S18). The predicted states of the toggle switch system, sampled every 25 minutes (8.4 samples within each of the forcing period of Exp5) in Fig. 11A, demonstrate that the kernel filter outperforms the particle filter. The trajectory length is 50 minutes (every second measurement), to increase the filter error and make a clear distinction between the two filters’ performances. This can be seen at each increase of LacI; the kernel filter has fitted the model parameters sufficiently, such that its LacI trajectories also increase. Whereas the particle filter trajectories immediately descend on all three examples. It can also be seen that at 300 minutes, the particle filter trajectory of LacI rapidly increases away from that of the experimental LacI concentration. The identification *MAE* (Eq.(7)) of the kernel filter, 0.3080, is less than the identification *MAE* of the particle filter, 0.3880.

**Figure 11.**
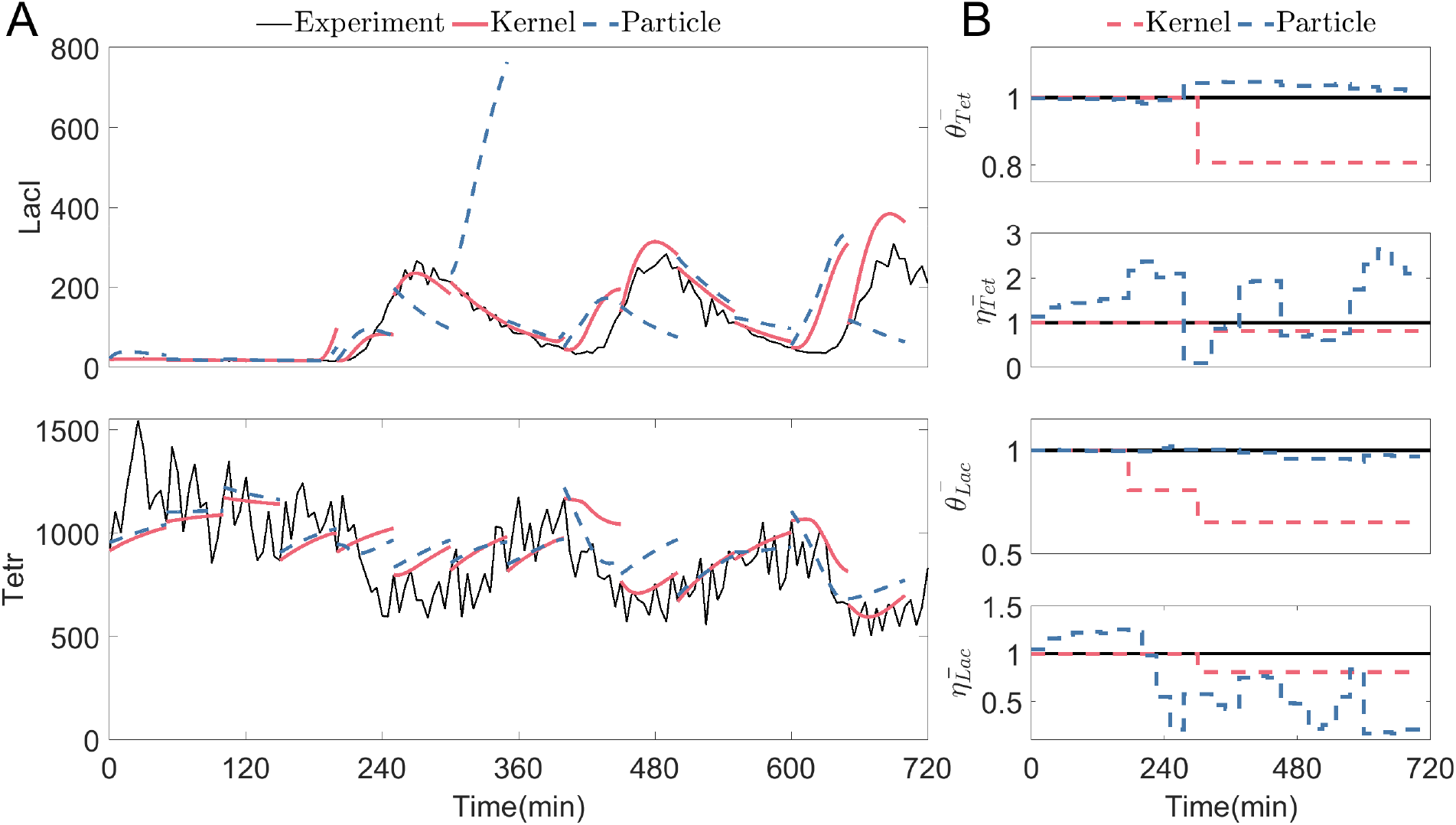
The filters fitting the parameters, online, to match the states of the toggle switch system, Eq.(17) to Eq.(22), to the states of the experimental data. A sample every 25 minutes is used, which is considered a low sampling regime here. Other simulation parameters: 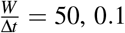 is the expected variance of parameters, *M* = 5000, 0.5 re-sampling threshold. A) Every 50 minutes, the prediction trajectory, 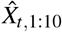 of each filter is compared to the experimental cell average (as seen in Fig. S18) using Exp5. B) The parameter fitting of 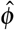, where each parameter has been normalised by *ϕ*_0_.

The corresponding parameter variations are shown in Fig. 11B. It can be seen that the particle filter changes the response of the model by altering both hills coefficients *η*_*Tet*_ and *η*_*Lac*_, with minimal changes in either *θ*_*Tet*_ or *θ*_*Lac*_. The kernel filter does the opposite, varying dissociation constants (*θ* s) more than the hills coefficients (*η*s). The Hill coefficients approximate the number of bound ligands between the two reactant states, whereas the dissociation constants are the rate of reaction. Therefore, the Hill coefficient usually tends towards integer values, showing that the Kernel filter parameter fittings make more biological sense, as the rate of reaction varies between cells and sample times, but the effect of the number of reactions within the system is fixed.

#### 3.3.2 Varied sampling rate identification

The identification Mean Absolute Error, *MAE* Eq.(7), indexing the error in the prediction trajectories (as in Fig. 11) is compared over a range of sampling rates in Fig. 12, matching the simulations in Fig. 5A and Fig. 9A. The experimental data collects samples every five minutes, therefore this is the fastest sampling rate used within these simulations, as this is the region of sample rates used within microfluidic devices. Various samples were not given to the filters to simulate slower sampling rates. For example, at a sampling rate of 10 minutes, every other sample is given to the filters.

**Figure 12.**
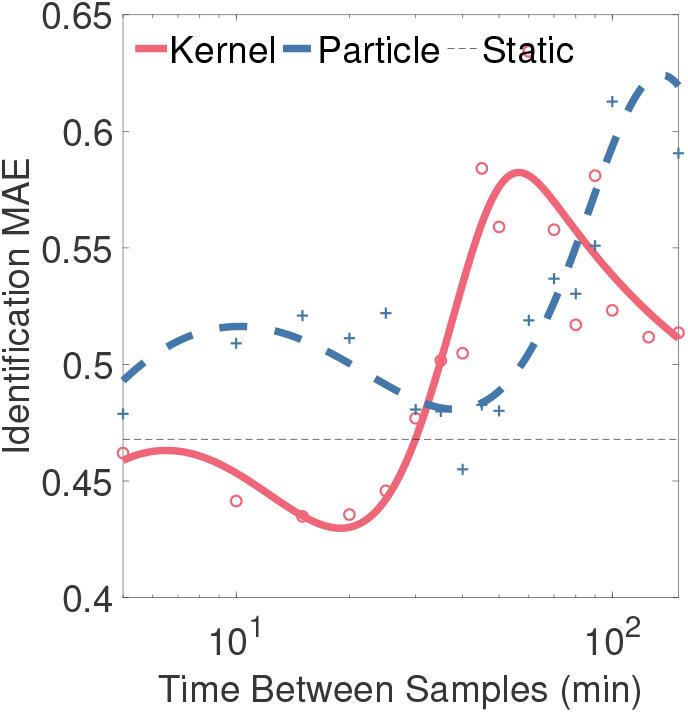
The identification *MAE*, Eq.(7), of the two filters is compared over a range of sampling rates for the toggle switch. The data points are the average identification *MAE* of 10 experiments (Exp2, Exp4, Exp5, Exp7, Exp8, Exp9, Exp10, Exp11, Exp12, Exp13, as discussed in Section S15) from [30]) at each time between samples. The simulation parameters for the kernel and particle filters are noted in Fig. 11.

Each cell within a microfluidics chip will have different rates of reaction, and therefore differ from the *in silico* model in Eq.(17) to Eq.(22), as can be seen in Fig. S14 to Fig. S27. Therefore, the static line in Fig. 12, is the mean of 100 simulations of the toggle switch model (Eq.(17) to Eq.(22)) where the parameters are selected from the random distributions described in [47], similarly based off the mean parameter values in [30], displaying the expected performance of a static model when used for state trajectories.

For the state trajectories plots of the synthetic gene oscillator and P53 oscillator in Fig. 5A and Fig. 9A, we describe three regions of the figure: a fast sampling region where both filters have a low identification MAE; a transition region, where the kernel filter retains a lower identification MAE than the particle filter; and finally a third region where both filter’s identification MAE tends to that of the static model. Due to the sample time limitations in the experimental data used in Fig. 12, we cannot see the first region, as the particle filter’s identification MAE has already tended to that of the static model. Therefore Fig. 12 starts within the transition region, where it can be seen that up until a sample time of 30 minutes (7 samples per period), the kernel filter tracking error remains significantly below that of the particle filter, showing that when sampling less than every 30 minutes, the kernel filter outperforms the particle filter. This transition region will be referred to as a low sample rate regime. In contrast, long sampling time, where neither filter outperforms the static parameters, is referred to as a very low sample rate regime. At sample times greater than every 30 minutes, the identification *MAE* of both filters tends to a random fluctuation near the static model line.

## 4 Conclusion

In this work, we demonstrated that kernel filters are valid for fitting model parameters according to online measurements at low sampling rates through simulations of three genetic networks. Particle filters are preferable for some plant systems at faster sampling rates, yet at reduced sampling rates, a kernel filter should be used (sampling less than once a minute for the synthetic gene oscillator, less than an hour for the P53 system and all experimentally viable sample rates of the toggle switch). Fitted mathematical models can then be added to a model-based controller to significantly increase controller performance at lower sampling rates.

The performance of the kernel filter could be improved in several ways. First, gradient-descent-based optimisers settle on local minima within the landscape of the *HSIC* cost function. A grid search algorithm has been used here, increasing the computational burden significantly with the number of parameters in the model and limiting the selected parameter values to defined discrete groups. A nonlinear solver that could find an acceptable minimum would drastically reduce the computational burden of adapting models with more fitted parameters. The kernel filter could also use Bayesian inference to find the optimal parameter set in the feature space (rather than the particle filter that uses Bayesian inference in the problem space).

Our results will facilitate the implementation of cybergenetics for multiple applications across biological systems, where practical considerations can significantly limit the sampling rate used for control.

## Acknowledgement

An EPSRC DTP Scholarship supports B.S. L.M. is supported by an EPSRC fellowship (grant EP/S01876X/1) and a BBSRC transition award (grant BB/W013959/1).

