## Supplementary Information for "Kernel Filter-Based Adaptive Controllers For Cybergenetics Applications"

### S1 Kernel Family

A kernel (also known as a covariance function) describes the transformation of a vector from a problem space to a feature space. The basis function used in this transformation can take many different forms.

The default choice is usually a Gaussian kernel, $\kappa_G$, [1]. With the hyperparameters, $\ell$ as the Gaussian length scale and $\sigma_n$ as the expected noise covariance that the kernel intrinsically adds to each measurement, determining how much the kernel trusts the underlying model.

$$\kappa_G = \exp\left(-\frac{(x-x')^2}{2\ell^2}\right) + \sigma_n \qquad \text{(S1)}$$

There are many different basis functions, as described in chapter four of [2] and [3]. Here, we compare seven commonly used kernel functions.

$$\kappa_{v=1/2} = exp\left(-\frac{(x-x')}{\ell}\right) + \sigma_n \qquad \text{(S2)}$$

The Matern kernels, $\kappa_v$, combine a polynomial with a Gaussian kernel, where $v$ determines the order of the polynomial.

$$\kappa_{v=3/2} = \left(1+\sqrt{3}\frac{(x-x')}{\ell}\right) exp\left(-\frac{\sqrt{3}(x-x')}{\ell}\right) + \sigma_n \qquad \text{(S3)}$$

$$\kappa_{v=5/2} = \left(1+\sqrt{5}\frac{(x-x')}{\ell} + \frac{5}{3}\frac{(x-x')^2}{\ell^2}\right) exp\left(-\frac{\sqrt{5}(x-x')}{\ell}\right) + \sigma_n \qquad \text{(S4)}$$

The rational kernel, $\kappa_R$, is equivalent to the summation of multiple Gaussian kernels with different length scales (as $\alpha \rightarrow$ inf, it forms a Gaussian kernel).

$$\kappa_R = \left(1+\frac{(x-x')^2}{2\alpha\ell^2}\right)^{\alpha} + \sigma_n \qquad \text{(S5)}$$

The periodic kernel, $\kappa_P$, includes a periodic exponent adapted to oscillation measurements where the hyperparameter $p$ denotes the expected period of the signal.

$$\kappa_P = exp\left(-\frac{2\sin^2(\pi|x-x'|/p)}{\ell/2}\right) + \sigma_n \qquad \text{(S6)}$$

The locally periodic kernel, $\kappa_{LP}$, combines the Gaussian kernel and a periodic kernel.

$$\kappa_{LP} = exp\left(-\frac{2\sin^2(\pi|x-x'|/p)}{\ell/2}\right) \exp\left(-\frac{(x-x')^2}{2\ell^2}\right) + \sigma_n \qquad \text{(S7)}$$

The ideal kernel family will be different depending on the problem. One hundred noisy simulations have been done, and one noise-free simulation for each of the seven kernel families for the P53 oscillator (Section 3.2), implementing the system identification problem discussed in Section 3.2.1. The resulting identification *MAE* (Eq. (7)) box plots are shown in Fig. S1. The hyperparameters are optimised online, as described in Section S2.

It was found that the locally periodic and Matern 3/2 kernels produced singular kernel matrices; therefore, the inverse needed for the hyperparameter optimisation could not be found. This was found for all initial hyperparameter estimations; thus, these kernels would not be chosen for this example and are not included in Fig. S1. It can be seen that Periodic, Matern 1/2 and Matern 5/2 kernels do significantly better than Gaussian and Rational kernels. The Periodic kernel has no outliers and a smaller range. The noise-free simulation is below any noisy simulations, indicating that the simulations would improve on average if the noise is reduced. Therefore, the periodic kernel has been used for all simulations within the kernel filter of the P53 oscillator. As the Synthetic gene oscillator (Section 3.1) is also periodic, and the toggle switch (Section 3.3) is forced to be periodic, it has also been decided to use a periodic kernel for all kernel filter applications within this report.

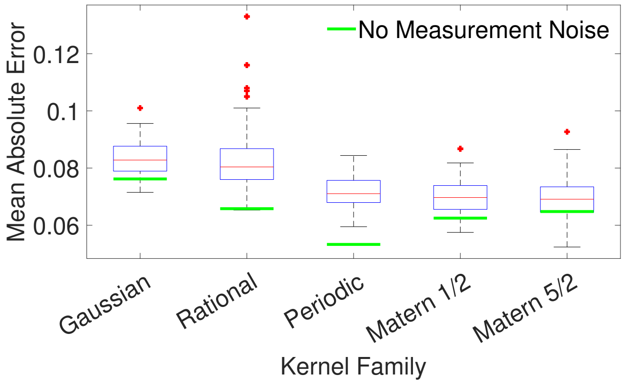


Figure S1: Comparison of the kernel families comparing 100 samples including 20dB noise to a noise-free simulation. Simulation parameters as described in Fig. 7.

### S2 Kernel Hyperparameter Optimisation

At each iteration of the kernel filter, the model kernel, $\kappa^x$, will vary with each trial parameter set, $\tilde{\phi}$, as described in Section 2.1. This means the optimisation would choose different hyperparameters for each $\tilde{\phi}$. If the model were an exact representation of the system, the model kernel, $\kappa^x$, would be identical to the measurement kernel, $\kappa^y$, which is fixed at each time step. Therefore, the kernel's hyperparameters are optimised over the fixed measurements for that step using $\kappa^y$. These same hyperparameters are used in the model kernels, $\kappa^x$.

The hyperparameters of the periodic kernel are $p$, $\ell$ and $\sigma_n$. The expected noise, $\sigma_n$, is assumed to be the same for all states. However, $p$ and $\ell$ are unique to each state. For a three state system, there are seven hyperparameters. These hyperparameters form a set, $\mathscr{H}$. The distribution of the measurement vector is dependent on the model parameters and kernel hyperparameters, $P(Y|\tilde{\phi},\mathscr{H})$.

The hyperparameter optimisation aims to find $\mathscr{H}$. We can use marginalisation to integrate out the hyperparameters and obtain the probability distribution of the outputs as,

$$P(Y|\tilde{\phi}) = \int P(Y|\tilde{\phi},\mathscr{H})P(\mathscr{H}|\tilde{\phi})d\mathscr{H}. \tag{S8}$$

For the particle filter, the marginalisation integral is undefined and therefore, marginal likelihood is used as an approximation [4]. However, for the kernel, the hyperparameters are all Gaussian processes of the form, $P(\mathscr{H}|\tilde{\phi}) \sim \mathscr{N}(0,\kappa)$, [2]. This means that the integral is tractable, resulting in the log marginal likelihood,

$$\log(P(Y|\tilde{\phi},\mathscr{H})) = -\frac{1}{2}Y^T\kappa^{-1}Y - \frac{1}{2}\log(|\kappa|) - \frac{\eta}{2}\log(2\pi), \tag{S9}$$

where $\eta$ is the number of states in the system.

We wish to optimise the hyperparameters to maximise the log marginal likelihood, $\max_{\mathscr{H}}(P(Y|\tilde{\phi},\mathscr{H}))$. Therefore, we use a gradient descent nonlinear optimiser (MATLAB's® fmincon) to find the local optimum from the initial conditions. We also provide the nonlinear optimiser with the partial differential of each hyperparameter, as in [2],

$$\begin{gathered}\frac{\partial}{\partial \mathscr{H}_j}\log(P(Y|\tilde{\phi},\mathscr{H})) = \frac{1}{2}tr\left((\alpha\alpha^T - \kappa^{-1})\frac{\partial\kappa}{\partial\mathscr{H}_j}\right)\\ \alpha = \kappa^{-1}Y\end{gathered} \tag{S10}$$

This optimisation of the hyperparameters is computationally heavy due to the need to calculate $\kappa^{-1}$, which can also lead to the singularity issues discussed in Section S1. The initial conditions for the simulation's hyperparameters depend on that simulation's sampling time, as discussed in Section S3. The optimised hyperparameters for the five plotted kernel simulations are shown in Fig. S2.

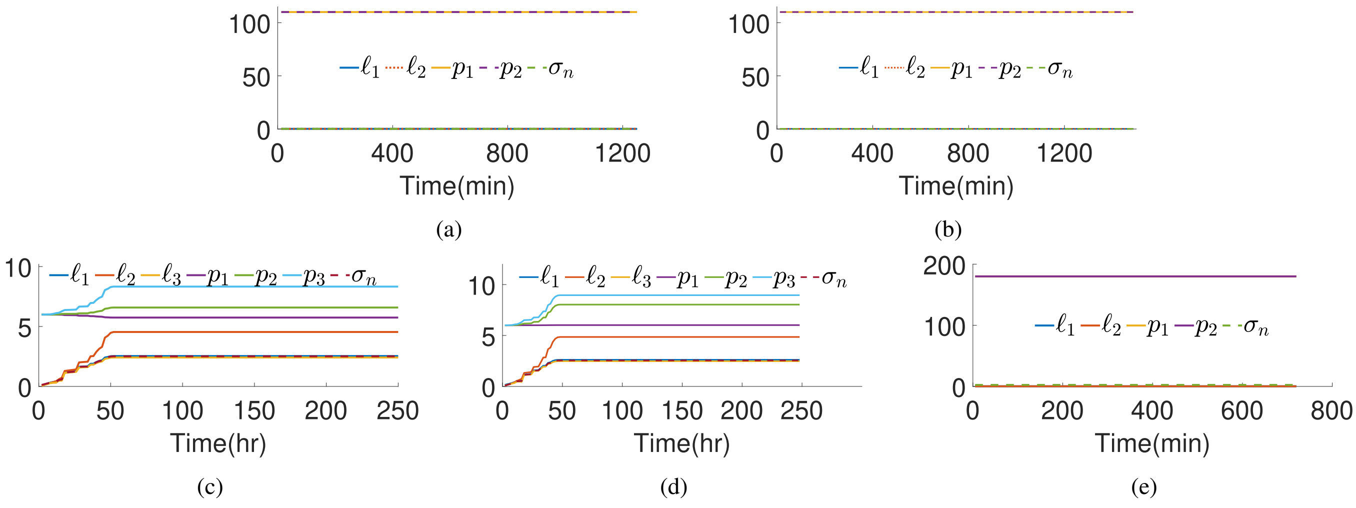


Figure S2: The optimisation of the kernel hyperparameters during the figures in the main text. a) Synthetic gene oscillator filtering, Fig. 3. b) Synthetic gene oscillator control, Fig. 4. c) P53 filtering, Fig. 7. d) P53 control, Fig. 8. e) Toggle switch filtering, Fig. 11.

### S3 Kernel Filter Parameter Sampling Rate Dependency

In a lab set-up, the measurement sampling time, $T_s$, will likely be fixed due to the equipment used. The filter/controller would be tuned to this specific case. However, for this paper, we are comparing the performance of the controller/filter at different sampling times. Table S1 presents some tuning guidelines to simplify the choice of the sampling time-dependent filter parameters at different sampling times.

#### S3.1 Parameter Search Mesh

As mentioned in Section 2.1, the kernel filter requires a grid search with a defined mesh to test each element of the set $\Phi_t$. For a system containing $n$ parameters, the discrete set, $\Phi_t$, contains all $3^n$ possible combinations of $\tilde{\phi}_t$ where each element of $\hat{\phi}_{t-1}$ has been multiplied by an element of $([1-\rho, 1, 1+\rho])$ to create a $3^n$ unique trial $\tilde{\phi}_t$.

It is assumed that the system parameters that make up $\phi$ vary continuously. If a constant mean and variation are assumed, the longer the time between the samples, the more the parameter could have changed since the last sample. Therefore, if the time between samples is larger (a slower sampling rate), the mesh size of the grid search should be larger. Based on numerical experimentation, we propose $\rho = KC_1\sqrt{T_s}$, where $KC_1$ is a tuned constant and $T_s$ is the time between samples. This rule was found to work well for the middle sections of Fig. 5, Fig. 9 and Fig. 12 (i.e. the time between samples shown as the middle condition in the second column of Table S1). However, it was seen that this rule breaks down at either end of the plot.

For the small sampling times (fast sampling rates), the filters observe the system frequently enough that the change in parameters is far less than the grid mesh. This means that the grid spacings can be very small, and the filter keeps up with the change. Therefore, a smaller mesh size can be used, and $KC_1$ is reduced for these faster sampling times, as shown in the second column of Table S1.

The continuous rule ($\rho = KC_1\sqrt{T_s}$) would result in a few large parameter jumps for the very large sampling time simulations (slow sampling rates), coupled with the controllers not receiving enough information to keep up with the system changes, results in large fluctuations around the static model line. These fluctuations are seemingly random. It has been decided that at these large sampling times, it would be better to make $\rho$ small once again so that the ill-informed parameter jumps are smaller and have less effect on the filter/control performance, therefore defining $\rho = KC_1$ where $KC_1$ is small.

Therefore, it was decided to split the mesh size $\rho$ into three steps. The middle step uses $KC_1\sqrt{T_s}$. Where either the identification *MAE* (Eq. (7)) or the control *RMSE* (Eq. (9)) fluctuates around a constant lower index value, the mesh size is reduced, $\rho = KC_1\sqrt{T_s}$. Where the identification *MAE* or control *RMSE* fluctuates around the static model line, the smaller mesh size is initiated, $\rho = KC_1$. Table S1 shows the specific constants and boundaries.

| Constant | $\rho(KC_1)$ | $KC_2$ | $KC_3$ | $KC_4$ |
|---|---|---|---|---|
| Value in synthetic gene oscillator identification - Fig. 3, 5A and S2a | $\rho = \begin{cases} 0.01\sqrt{T_s}, & \text{if } T_s < 2 \\ 0.04\sqrt{T_s}, & \text{if } 2 \leq T_s \leq 50 \\ 0.001, & \text{if } T_s > 50 \end{cases}$ | $10^{-12}$ | 0.001 | 0.001 |
| Value in synthetic gene oscillator control - Fig. 4, 5B, C and S2b | $\rho = \begin{cases} 0.01\sqrt{T_s}, & \text{for all } T_s \end{cases}$ | $10^{-12}$ | 0.001 | 0.001 |
| Value in P53 identification - Fig. 7, 9A and S2c | $\rho = \begin{cases} 0.005\sqrt{T_s}, & \text{if } T_s < 0.5 \\ 0.01\sqrt{T_s}, & \text{if } 0.5 \leq T_s \leq 8 \\ 0.01, & \text{if } T_s > 8 \end{cases}$ | $10^{-3}$ | 0.001 | 0.001 |
| Value in P53 control - Fig. 8, 9B, C and S2d | $\rho = \begin{cases} 0.005\sqrt{T_s}, & \text{if } T_s < 0.09 \\ 0.01\sqrt{T_s}, & \text{if } 0.09 \leq T_s \leq 10 \\ 0.01, & \text{if } T_s > 10 \end{cases}$ | $10^{-3}$ | 0.001 | 0.001 |
| Value in toggle switch identification - Fig. 11, 12, S2e and S30 | $\rho = \begin{cases} 0.00005\sqrt{T_s}, & \text{if } T_s < 5 \\ 0.005\sqrt{T_s}, & \text{if } 5 \leq T_s \leq 25 \\ 0.0005, & \text{if } T_s > 25 \end{cases}$ | $10^{-3}$ | 0.0001 | 0.0001 |

Table S1: A guideline for the kernel filter parameters that are dependent on sampling time: $\rho(KC_1)$ dictates the sample grid spacing; $KC_2$ dictates the hyperparameter variance used within the hyperparameter optimisation; $KC_3$ dictates the initial estimation for the hyperparameters, $\ell$; and $KC_4$ dictates the initial estimation for the hyperparameters, $\sigma_n$.

#### S3.2 Hyperparameter Variance

Similarly to $\rho$ for the system parameter changes, the kernel hyperparameters will also change over time. The larger the sampling times (slower sampling rate), the larger the temporal gap between the rows/columns of the kernel, and therefore, there will be a larger variance associated with the hyperparameters. The continuous rule has been set to $KC_2T_s^2$. The origin of the square relationship comes from the Gaussian nature of the kernel basis functions. The constant for each simulation can be seen in Table S1.

#### S3.3 Hyperparameter Initial Estimation

The kernel hyperparameters are not unknown covariances (such as the tuning of the particle filter); they have a somewhat physical representation dependent on the time signal from which the kernel is produced. $p_i$ is the period of state $i$ and, therefore, does not change with the sampling time of the response and is independent of the sampling time. For all simulations, $p_i$ is initially set to the free response period or the period of the data's inputs. The Gaussian length, $\ell$, dictates the expected difference between observations. Therefore, the smaller the sampling times, the smaller the expected difference between the measurements (as long as the sampling time is much less than the time period). Therefore, $\ell$ is also expected to vary with the sampling time. The initial estimate of $\ell$ at the beginning of the simulations is given by the continuous rule $KC_3T_s^2$. Finally, the expected noise in the states, $\sigma_n$, will also vary with the sampling time, as it is proportional to the states. Therefore, a similar continuous rule is used, $KC_4T_s^2$. $\sigma_n$ denotes how much the kernel trusts the underlying model of the system and will thus still be non-zero for noise-free simulations, as the model is imperfect. The constant for each simulation can be seen in Table S1.

### S4 Particle Filter Parameter Sampling Rate Dependency

#### S4.1 Particle Variance

To compare the two filter types, there needs to be some consistency in how the filters are tuned. Each filter implements a different search method for the parameter set, $\phi$. The kernel filter uses the grid search, and the particle filter uses Bayesian inference. The spacing of the grid, $\rho$, is a discrete representation of the expected change in the parameters per time step (for a discrete grid search), whilst the particle filter's variance, $\sigma$, is a continuous representation of the expected change in the parameters, per time step, in this continuous search space. Both $\rho$ and $\sigma$ fundamentally describe the expected change in each parameter, and therefore, the same tuning rules are used.

The variance, $\sigma_i$ of each parameter, $i$ (of a total $n$), within the particle sets, $\tilde{\phi}$ follows the continuous rule, $\sigma_i = PC_i\sqrt{T_s}$. The same observations about the larger and smaller sampling times (as discussed in Section S3.1) result in the same three-stepped tuning. Where either the identification $MAE$ (Eq. (7)) or the control $RMSE$ (Eq. (9)) begins to fluctuate around a constant lower index, the mesh size is also reduced, $\sigma_i = PC_i\sqrt{T_s}$. Where the index fluctuates around the static model line, the smaller mesh size is initiated, using $\sigma_i PC_i$. Table S2 shows the specific constants and boundaries.

| Constant | $[\sigma_a(PC_a), \sigma_b(PC_b)]$ | $PC_{n+a}$ | $PC_{n+b}$ |
|---|---|---|---|
| Value in synthetic gene oscillator identification Fig. 3 and 5A | $[\sigma_a, \sigma_b] = \begin{cases} \left[10^{-1.5}, 10^{-3}\right]\sqrt{T_s}, & \text{for all } T_s \end{cases}$ | $10^{-2}$ | $10^{-3}$ |
| Value in synthetic gene oscillator control Fig. 4, 5B and D | $[\sigma_a, \sigma_b] = \begin{cases} \left[10^{-1.5}, 10^{-3}\right]\sqrt{T_s}, & \text{if } T_s < 2 \\ \left[10^{-2.5}, 10^{-3}\right]\sqrt{T_s}, & \text{if } 2 \le T_s \le 5 \\ \left[10^{-1.5}, 10^{-3}\right], & \text{if } T_s > 5 \end{cases}$ | $10^{-2}$ | $10^{-3}$ |
| Value in P53 identification Fig. 7 and 9A | $\sigma_a = \begin{cases} 10^{-5}\sqrt{T_s}, & \text{if } T_s < 0.1 \\ 10^{-4}\sqrt{T_s}, & \text{if } 0.1 \le T_s \le 2 \\ 10^{-5}, & \text{if } T_s > 2 \end{cases}$ | $10^{-3}$ | |
| Value in P53 control Fig. 8, 9B and D | $\sigma_a = \begin{cases} 10^{-4}\sqrt{T_s}, & \text{if } T_s < 1 \\ 10^{-4}\sqrt{T_s}, & \text{if } 1 \le T_s \le 10 \\ 10^{-6}, & \text{if } T_s > 10 \end{cases}$ | $10^{-6}$ | |
| Value in toggle switch identification Fig. 11, 12 and S30 | $\sigma_a = \begin{cases} 0.0005\sqrt{T_s}, & \text{if } T_s < 5 \\ 0.005\sqrt{T_s}, & \text{if } 5 \le T_s \le 25 \\ 0.0005, & \text{if } T_s > 25 \end{cases}$ | $10^{-1}$ | |

Table S2: A guideline for the particle filter parameters that are dependent on sampling time: $\sigma_a(PC_a)$ and $\sigma_b(PC_b)$ dictate the variance of each element of the particle; and $PC_{n+a}$ and $PC_{n+b}$ dictates the initial spread of each element of the particle. For the synthetic gene oscillator (Section 3.1), $a = 1$ and $b = 2$. For the P53 system (Section 3.2) $a$ contains all three of the fitted parameters, $a = [2, 3, 6]$. For the toggle switch (Section 3.3) $a$ contains all four of the fitted parameters.

#### S4.2 Particle Initial Spread

The particle filter requires the user to define the initial spread of particles. The particles will be re-sampled during the simulation if they coalesce on one set. Therefore, we only need to define the initial spread of the particles. Crucial to the first step of the particle filter, ensuring that $\phi_1$ is within the spread. As discussed in Section S3.1, with the random walk, the longer the time between samples, the further $\phi_1$ is likely to be from $\phi_0$, and therefore, the larger the initial spread should be. Similarly to $\rho$ and $\sigma_i$, the continuous rule $PC_{3n+i}\sqrt{T_s}$ is used. The values of these constants can be seen in Table S2.

### S5 Synthetic Gene Oscillator

The model of synthetic gene oscillator found in [5] describes the interactions between a CI protein ($X_1$) and a LAC protein ($X_2$). This paper discusses the model order reduction of a four state kinetics model, containing both the proteins and their corresponding dimers, to the two state model containing a dimensionless representation of the CI protein as $x_1$ and a dimensionless representation of the LAC protein as $x_2$, as shown in Eq. (10) and Eq. (11). The CI protein promotes both the CI and LAC dimers, whereas the LAC protein inhibits both dimers, as shown in Fig. S3.

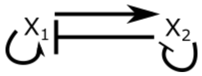


Figure S3: Synthetic gene oscillator.

#### S5.1 Parameters in the Synthetic Gene Oscillator Model

We define the state vector, $\mathbf{x} = [x_1, x_2]^T$. The dimensionless initial conditions $\mathbf{x}_0 = [0.3, 2]^T$ are used in all simulations. The model parameters are described in Table S3.

| Variable | Definition | Value [5] |
|---|---|---|
| $\alpha$ | Summarises the production rates from the plasmids. The rate of transcription/translation resulting in the production of both proteins. | 11 |
| $\sigma$ | Summarises the rate of the protein reactions. It is the ratio of the binding rate of the dimer to the production rate of the dimer protein complex for both proteins. | 2 |
| $\tau$ | The time scale separation of the two protein states. It is physically the ratio of the copy numbers of the two plasmids. | 5 |
| $u_{x_1}$ | The degradation rate of the CI protein. Used as an input in the control system as the degradation of the CI857 protein is tunable by temperature. | 0.105 $min^{-1}$ (free response) |
| $u_{x_2}$ | The degradation rate of the LAC protein. Used as an input in the control system as isotropyl-b-D-thiogalactopyranoside (IPTG) can be added to increase the degradation. | 0.036 $min^{-1}$ (free response) |

Table S3: Parameters used in the synthetic gene oscillator model, including the two input parameters.

### S6 Synthetic Gene Oscillator with Noisy Measurements

The simulations in the main text do not include measurement noise to better visually demonstrate the superior performance of the kernel filter at low sampling rates. However, this is not realistic for actual experiments. Therefore, noisy simulations of both the filtering and control simulations are displayed in Fig. S4 and Fig. S5 for the synthetic gene oscillator. The measurement noise includes a covariance $\xi = 10^{-2.5}$, resulting in a root mean squared, signal-to-noise ratio of approximately $20dB$.

For the same magnitude of measurement noise, the filtering simulations in Fig. S4 show that the prediction horizon of the kernel filter still outperforms the particle filter (identification $MAE$ - 0.0758 $<$ 0.1950). The plotted state is not measured and, therefore, is not masked by the noise, demonstrating that even in the presence of noise, the kernel outperforms the particle filter at low sampling rates.

It can be seen in Fig. S5 that as the actual output is displayed, the performance of the controllers is masked by the random noise added to the output measurement. This complicates the visual comparison between the filters' performance. However, the control $RMSE$ for the kernel filter is still less than the particle filter (control $RMSE$ - 0.1718 $<$ 0.2219).

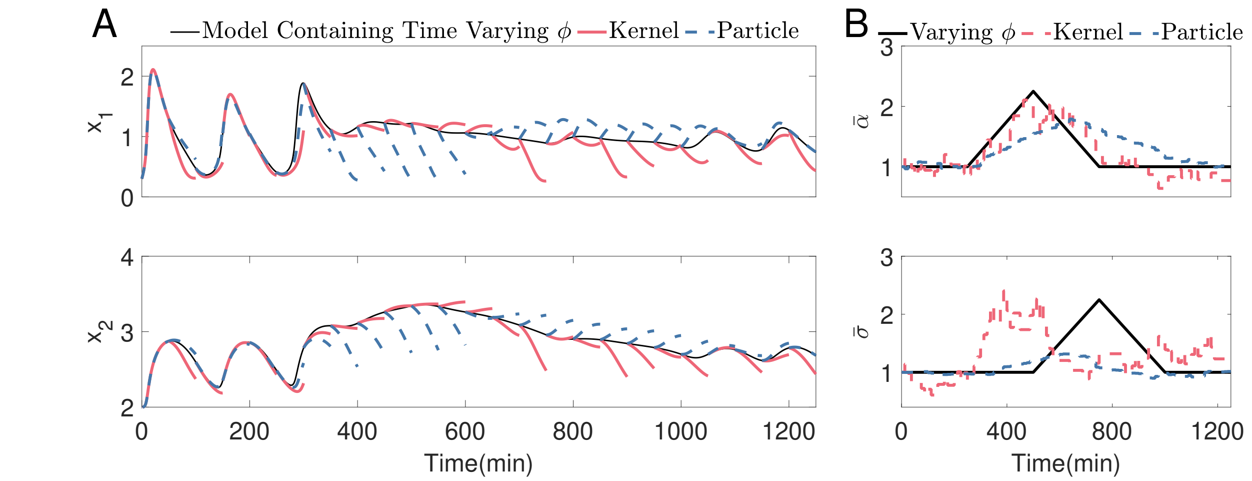


Figure S4: Low sampling rate filtering for the Synthetic gene oscillator in Section 3.1, including measurement noise. a) The kernel and particle filters fitting the parameters online to match the states of a system in which the true parameter value, $\phi$, changes over time. Every 50 minutes, the prediction whiskers are recentered to the corresponding measurement, $Y_t$, and the predictive model, $\hat{X}_{t:t+50}$, is plotted. The Filters are used to estimate the nonlinear parameter set used within the model (Eq.(10) - Eq.(11)). One sample every 12.5 minutes (8.8 samples per period) is used, which is considered a low sampling regime here. b) The corresponding parameter changes. Other simulation parameters: $\frac{W}{\Delta t} = 500$, $M = 5000$, 0.5 re-sampling threshold, 0.0001 expected variance of parameters.

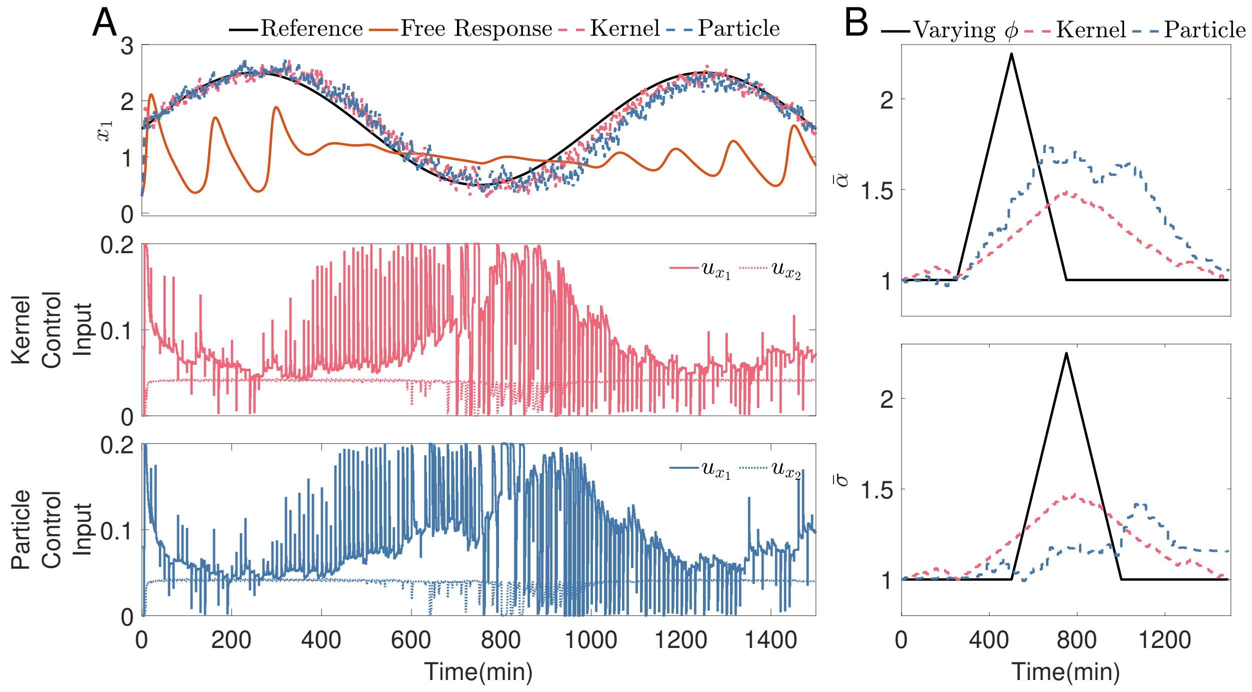


Figure S5: Low sampling rate filtering and control for the Synthetic gene oscillator in Section 3.1, including measurement noise. a) The system's output qualitatively follows the sine wave reference using the control input shown. The filters fitting the parameters online to match the states of the synthetic gene oscillator in which the nominal parameter value, $\phi$, changes over time. One sample every 12.5 minutes (8.8 samples per period) is used, which is considered a low sampling regime here. b) The corresponding parameter changes. Other simulation parameters: the control input is actuated once a minute; $N = 10$ minutes, containing ten input steps $\frac{W}{\Delta t} = 50$, 0.0001 is the expected variance of parameters, $M = 5000$, 0.5 re-sampling threshold.

### S7 P53 Oscillator

The model of a P53 oscillator describes the interactions between the p53 protein and the Mdm2 protein. Model 4 from [6] is used here, containing three states: p53 protein as $x_1$; the Mdm2 precursor as $x_2$ and the Mdm2 protein as $x_3$, as shown in Eq. (13) to Eq. (15). The model is based on mass action kinetics, containing a basal production of P53, triggering a chain reaction to produce Mdm2 through its precursor. The system includes its own feedback loop through a Michaelis Menten term, describing the inhibition of P53 from Mdm2. The input to the system is an extra degradation of Mdm2 via the addition of Nutlin, $u$ [7], a competitive inhibitor to P53. A schematic of the P53-Mdm2 interaction is shown in Fig. S6.

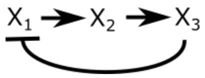


Figure S6: P53 oscillator.

#### S7.1 Parameters in the P53 Model

As above the state vector, $\mathbf{x} = [x_1, x_2, x_3]^T$. The initial conditions $\mathbf{x}_0 = [0, 0.1, 0.8]^T$ are used in all simulations. The model parameters are described in Table S4.

| Variable | Definition | Value |
|---|---|---|
| $\alpha_y$ | Mdm2 degradation rate. | $0.8h^{-1}$ [6] |
| $\alpha_0$ | Mdm2 maturation rate. | $0.9h^{-1}$[6] |
| $\alpha_k$ | Saturating p53 degradation rate. | $1.7P_{max}M_{max}^{-1}h^{-1}$ [6] |
| $k$ | p53 threshold for degradation by Mdm2. | $0.001P_{max}$ [6] |
| $\beta_x$ | p53 production rate. | $0.9P_{max}h^{-1}$ [6] |
| $\beta_y$ | p53-dependent Mdm2 production rate. | $1.2\ M_{max}h^{-1}$ [6] |
| $k_m$ | Nutlin rate constant | $200\ min^{-1}$ [7] |

Table S4: Parameters used in the P53 model including the input parameter.

### S8 P53 Oscillator with Noisy Measurements

As stated in Section S6, the simulations in the main text do not include measurement noise to better visually demonstrate the superior performance of the kernel filter at low sampling rates. This is not realistic for actual experiments; therefore, noisy simulations of both the filtering and control simulations are displayed in Fig. S7 and Fig. S8 for the P53 oscillator. The measurement noise includes a covariance $\xi = 10^{-2.5}$, resulting in a root mean squared, signal-to-noise ratio of approximately $20dB$.

For the same magnitude of measurement noise, the filtering simulations in S7 show that the prediction horizon of the kernel filter still outperforms the particle filter (identification $MAE$ - $0.0960 < 0.1650$). This plotted state is not measured and, therefore, is not masked by the noise, demonstrating that even in the presence of noise, the kernel outperforms the particle filter at low sampling rates.

It can be seen in Fig. S8 that as the actual output is displayed, the performance of the controllers is masked by the random noise added to the output measurement. This does not make an easy visual comparison between the filters. However, the control $RMSE$ for the kernel filter is still less than the particle filter (control $RMSE$ - $0.3600 < 0.4200$).

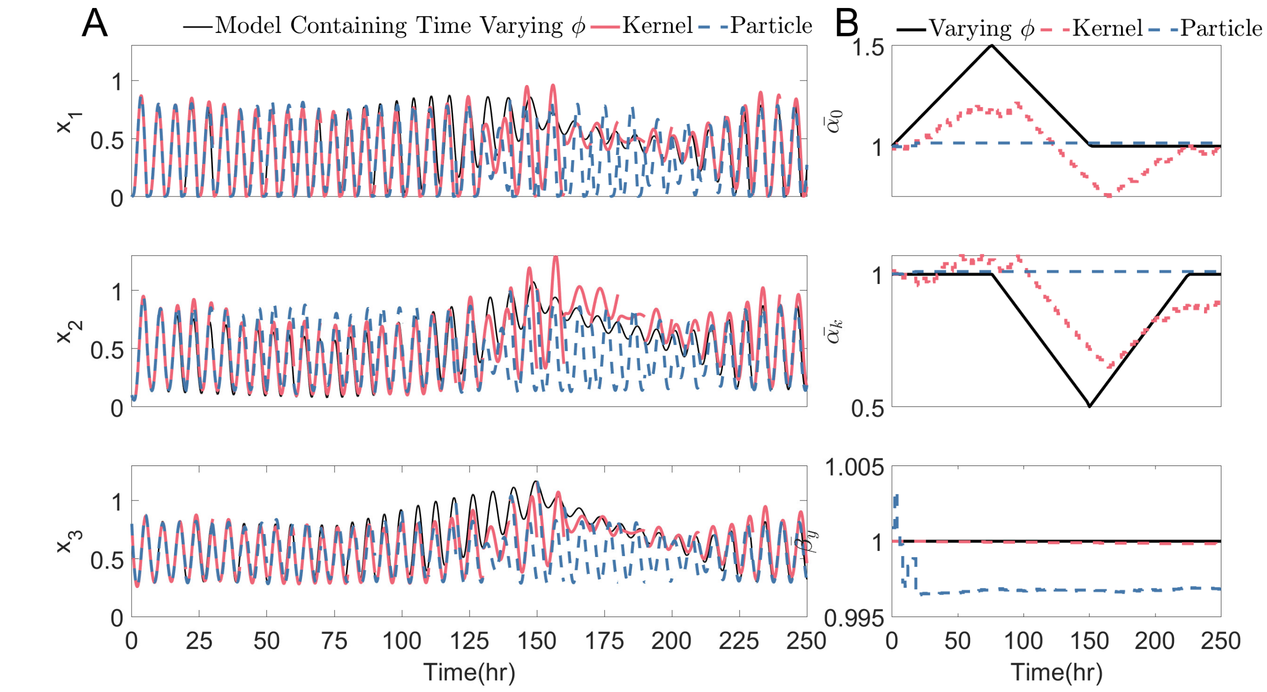


Figure S7: Low sampling rate filtering for the P53 oscillator, as in Section 3.2, including measurement noise, $\xi = 10^{-2.5}$. a) The kernel and particle filters fitting the parameters online to match the states of a system in which the true parameter value, $\phi$, changes over time. b) The corresponding parameter changes. Other simulation parameters: $\frac{W}{\Delta t} = 50$, $M = 5000$, 0.5 re-sampling threshold, 0.0158 expected variance of parameters.

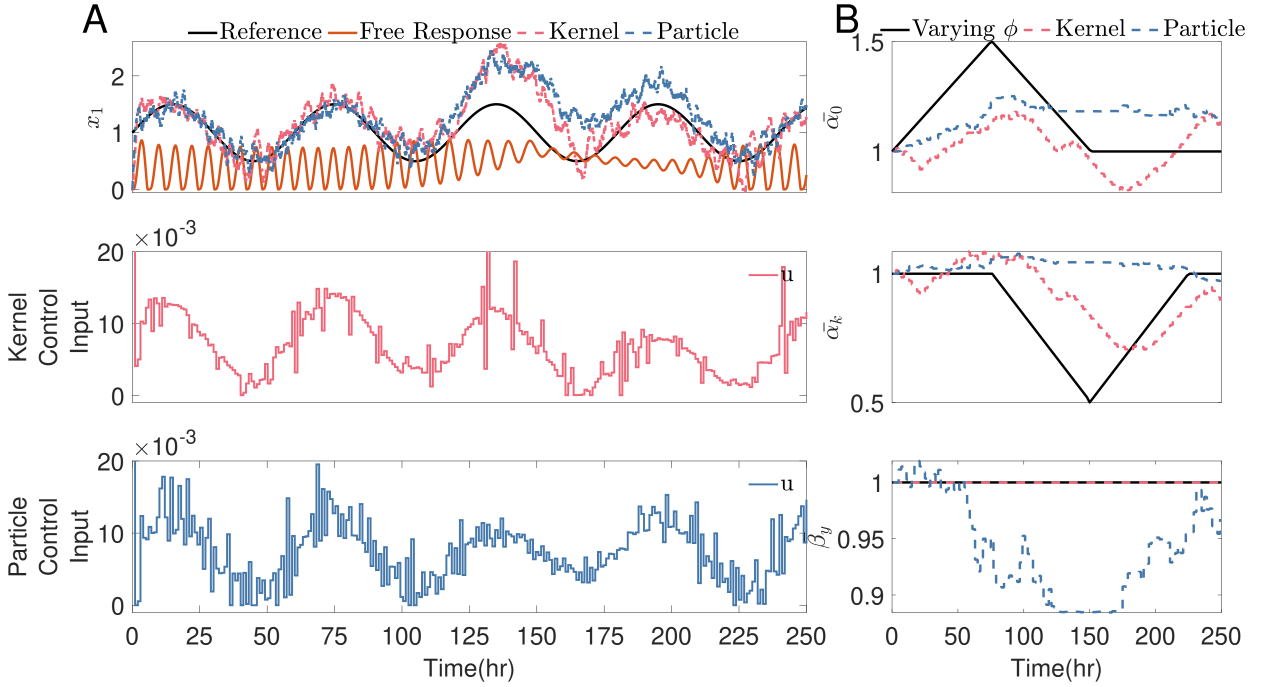


Figure S8: Low sampling rate control for the P53 oscillator, as in Section 3.2, including measurement noise, $\xi = 10^{-2.5}$. a) The system's output qualitatively follows the sine wave reference using the control input shown. The filters fit the parameters online to match the states of the P53 system in which the nominal parameter value, $\phi$, changes over time. b) The corresponding parameter changes. Other simulation parameters: the control input is actuated once an hour; $N = 10$ hours, containing two input steps, $\frac{W}{\Delta t} = 50$, 0.0158 is the expected variance of parameters, $M = 5000$, 0.5 re-sampling threshold.

#### S9 Long Sampling Times

Fig. S9 shows the performance of the kernel filter compared to the particle filter over a range of prediction horizons and sampling times for the P53 oscillator. The sample times included here far exceed those of Fig. 9, containing sample times that would not be used in-vitro, but are included here for discussion. The simulation is as described in Section 3.2.3, including the longer sampling times. It can be seen that there is a darker region for the kernel filter compared to the particle filter between sampling times of 6-40 hours (1-0.15 samples per period) and any prediction horizon, $N \geq 2$. It shows that not only when $N = 10$ (as in Fig. 9B) but for $N \geq 2$, the kernel filter outperforms the particle filter, except for a few 'light spots'.

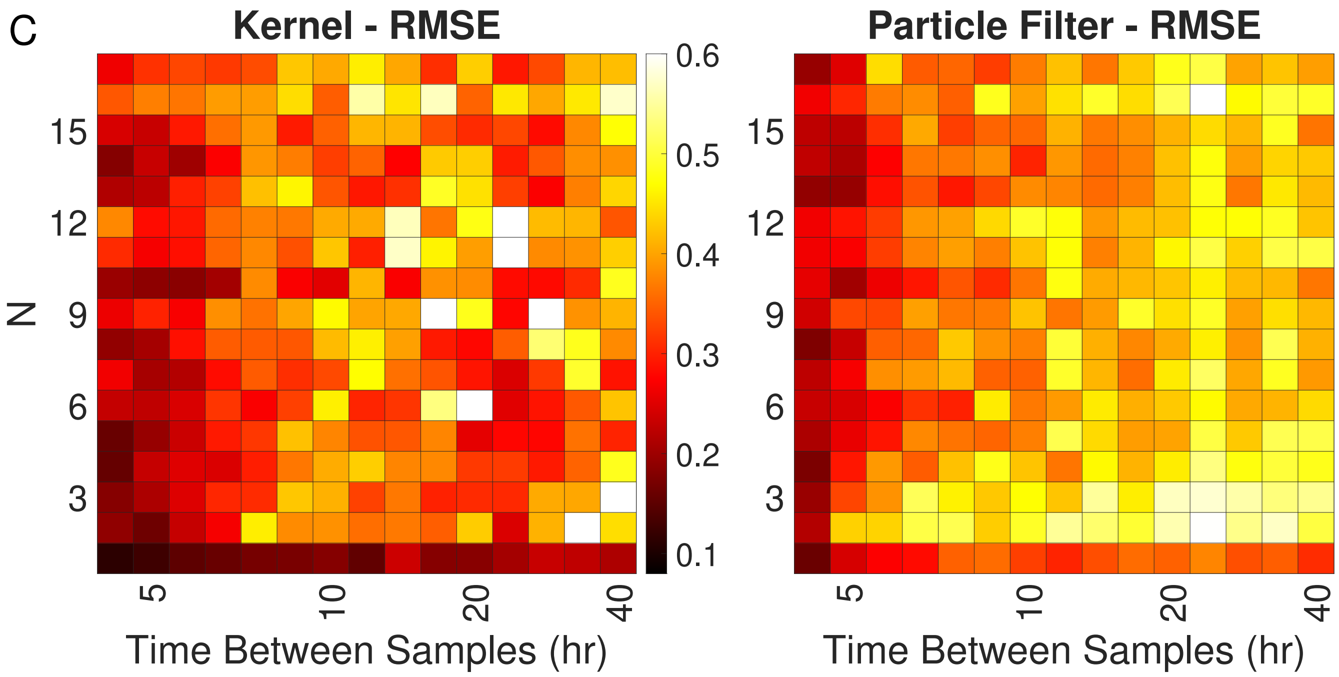


Figure S9: The control *RMSE* of the two filters, compared over a range of sampling rates and prediction horizons, $N$, for the P53 oscillator. Other simulation parameters as in Fig 9.

The particle filter can jump larger increments in $\hat{\phi}$, whereas the kernel filter is restricted to predetermined parameter jumps (as described in Section 2.1). As the non-linearities hide some of the changes in parameters from the filters, the particle filter can respond faster when it realises that it has been pushing $\hat{\phi}$ away from $\phi$. In contrast, the kernel filter can only take small predetermined steps back towards $\phi$. These light spots are simulations where the filter has initially strayed too far from $\phi$ to track anything like $X(\phi)$ as the simulation progresses. In the future, this could be improved for the kernel filter by improving the nonlinear optimisation described in Section 2.1.

### S10 P53 System With Six fitted Parameters

The P53 system (Eq.(13) - Eq.(15)) has six kinematic constants, yet for section 3.2 the two filters only vary $\alpha_0$, $\alpha_k$ and $\beta_y$. The identification and control simulations are repeated here, allowing the filters to vary all six parameters.

#### S10.1 Kernel and Particle Filter Simulation Parameters

| Constant | $\rho(KC_1)$ | $KC_2$ | $KC_3$ | $KC_4$ |
|---|---|---|---|---|
| Value in P53 identification - Fig. S10 and S12A | $\rho = \begin{cases} 0.005\sqrt{T_s}, & \text{if } T_s < 0.5 \\ 0.01\sqrt{T_s}, & \text{if } 0.5 \leq T_s \leq 8 \\ 0.01, & \text{if } T_s > 8 \end{cases}$ | $10^{-6}$ | 0.001 | 0.001 |
| Value in P53 control - Fig. S11, S12B and C | $\rho = \begin{cases} 0.005\sqrt{T_s}, & \text{if } T_s < 0.5 \\ 0.01\sqrt{T_s}, & \text{if } 0.5 \leq T_s \leq 8 \\ 0.01, & \text{if } T_s > 8 \end{cases}$ | $10^{-6}$ | 0.001 | 0.001 |

Table S5: A guideline for the kernel filter parameters that are dependent on sampling time: $\rho(KC_1)$ dictates the sample grid spacing; $KC_2$ dictates the hyperparameter variance used within the hyperparameter optimisation; $KC_3$ dictates the initial estimation for the hyperparameters, $\ell$; and $KC_4$ dictates the initial estimation for the hyperparameters, $\sigma_n$.

| Constant | $\sigma_a(PC_a)$ | $\sigma_b(PC_b)$ | $PC_{n+a}$ | $PC_{n+b}$ |
|---|---|---|---|---|
| Value in P53 identification Fig. S10 and S12A | $\sigma_a = \begin{cases} 10^{-5}\sqrt{T_s}, & \text{if } T_s < 0.1 \\ 10^{-4}\sqrt{T_s}, & \text{if } 0.1 \leq T_s \leq 2 \\ 10^{-5}, & \text{if } T_s > 2 \end{cases}$ | $\sigma_b = \begin{cases} 10^{-9}\sqrt{T_s}, & \text{if } T_s < 0.1 \\ 10^{-8}\sqrt{T_s}, & \text{if } 0.1 \leq T_s \leq 2 \\ 10^{-9}, & \text{if } T_s > 2 \end{cases}$ | $10^{-3}$ | $10^{-7}$ |
| Value in P53 control Fig. S11, S12B and C | $\sigma_a = \begin{cases} 10^{-3}\sqrt{T_s}, & \text{if } T_s < 0.5 \\ 10^{-4}\sqrt{T_s}, & \text{if } 0.5 \leq T_s \leq 1.2 \\ 10^{-5}, & \text{if } T_s > 1.2 \end{cases}$ | $\sigma_b = \begin{cases} 10^{-5}\sqrt{T_s}, & \text{if } T_s < 0.5 \\ 10^{-7}\sqrt{T_s}, & \text{if } 0.5 \leq T_s \leq 1.2 \\ 10^{-8}, & \text{if } T_s > 1.2 \end{cases}$ | $10^{-2}$ | $10^{-5}$ |

Table S6: A guideline for the particle filter parameters that are dependent on sampling time: $\sigma_a(PC_a)$ and $\sigma_b(PC_b)$ dictate the variance of each element of the particle; and $PC_{n+a}$ and $PC_{n+b}$ dictates the initial spread of each element of the particle. For the P53 system (Section 3.2), $a = [1,2,3,5,6]$ and $b = 4$.

#### S10.2 Low Sampling Rate Filtering

The parameter changes are identical to those discussed in Section 3.2.1. The trajectories are 10 hours long.

The predicted states of the P53 system, Fig. S10A, sampled every 2.5 hours (2.4 samples per period), demonstrate the kernel filter outperforming the particle filter. This can be seen between 25 and 100 hours with the decrease and then increase in amplitude of $x_2$ and between 125 and 225 hours, where neither filter perfectly matches the changes in the centre of the oscillation for all three states. Still, the kernel filter does qualitatively and quantitatively better. It can later be seen that this difference is significant in the control simulations between these times. The identification $MAE$ (Eq.(7)) of the kernel filter, 0.0667, is significantly less than the identification $MAE$ of the particle filter, 0.1600.

The corresponding parameter variations are shown in Fig. S10B. The particle filter fails to track any meaningful change in the states, whilst the parameters randomly vary around the initial value. The kernel filter varies all parameters; however, the change in $\hat{\phi}$ may be away from the 'true' parameter value, $\phi$. Like so many biological models, this model is over-parametrised, and non-identifiable [8], but can still achieve a reasonable state estimation. For instance, the increase in $\bar{\beta}_x$ (the production of $x_1$) rather than a decrease in $\alpha_k$ (the degradation of $x_1$), resulting in an equivalent sum change of $x_1$ over time, resulting in a good state prediction.

#### S10.3 Low Sampling Rate Control

The same reference as discussed in Section 3.2.2 is used here, and the performance for both filters can be seen in Fig. S11A for the P53 system, sampling every 2.5 hours (2.4 samples per period). Between 125 and 225 hours, the parameter changes cause the free response to change its centre of oscillation and amplitude. Both filters try to fit their models to keep up with this, but it can be seen that the particle filter model does not keep up with the changes, causing a large error from the reference between these times. The kernel's control $RMSE$ of 0.152 is less than the particle filter's control $RMSE$ of 0.274.

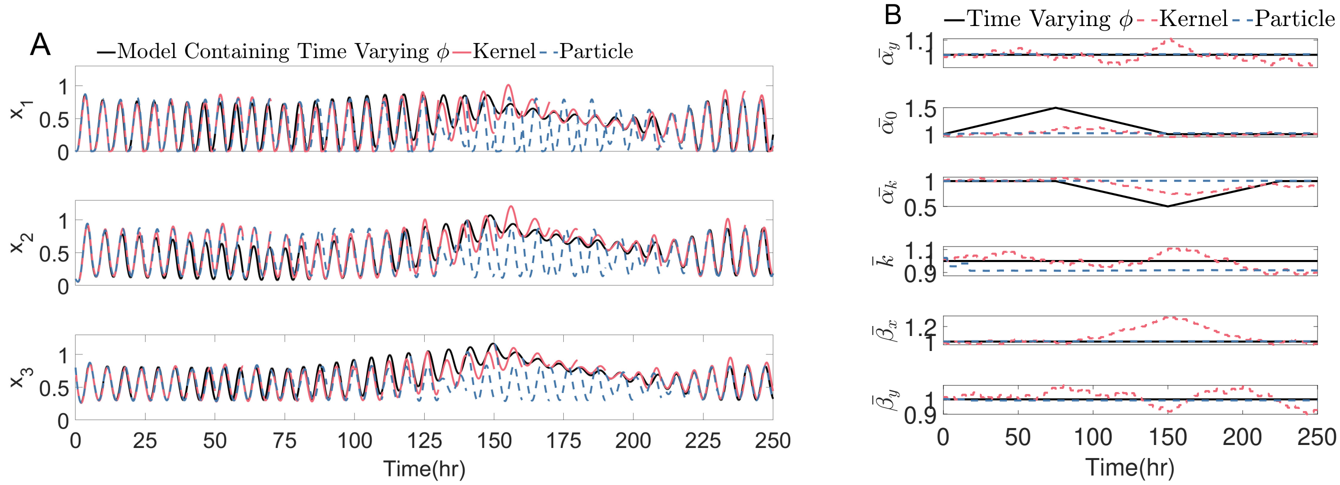


Figure S10: The filters fitting the parameters, online, to match the states of the P53 system (Eq.(13) - Eq.(15)) in which the true parameter value, $\phi$, changes over time. A sample every 2.5 hours (2.4 samples per period) is used, which is considered a low sampling regime here. Other simulation parameters: $\frac{W}{\Delta t} = 50$, 0.0158 is the expected variance of parameters, $M = 5000$, 0.5 re-sampling threshold. a) Every 10 hours, the prediction trajectory, $\hat{X}_{t:t+10}$. b) The parameter fitting of $\hat{\phi}$, compared to the true parameters, $\phi$. Each parameter has been normalised by the initial parameter estimation in Section 3.2.

The parameter changes within Fig. S11B for the P53 system are very similar to Fig S10B where the kernel filter does not follow the nominal parameter changes, $\phi$, but finds an alternative combination of the six parameters that produce a similar state response, as can be seen again with the increase in $\beta_x$ instead of a decrease in $\alpha_k$. The particle filter does not have an informed variation from the initial estimation at this sampling rate.

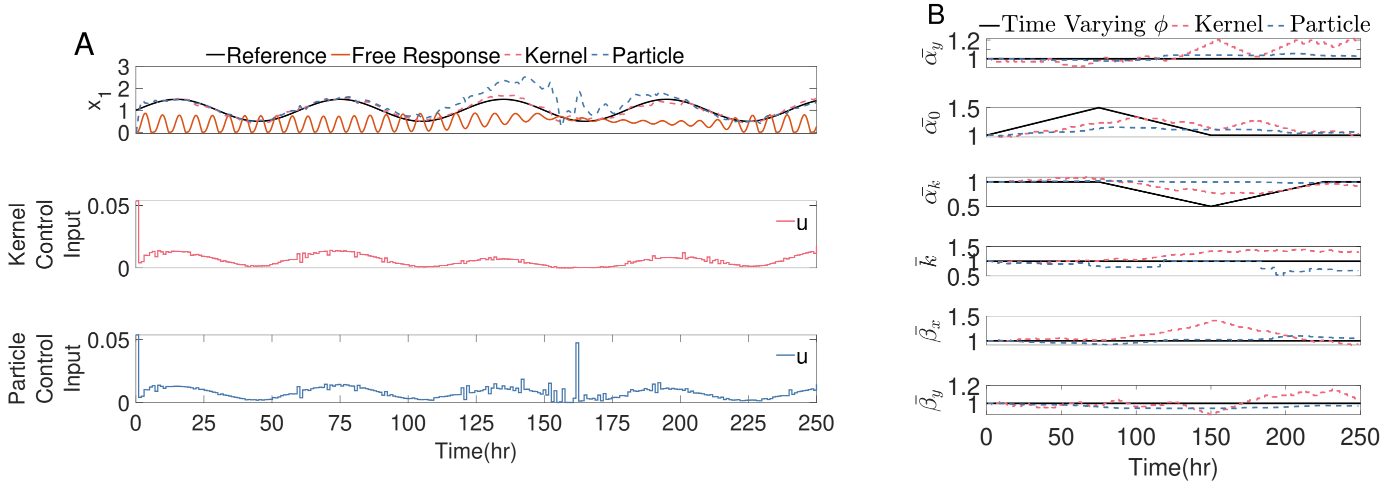


Figure S11: The filters fitting the parameters, online, to match the states of the P53 system (Eq.(13) - Eq.(15)) in which the true parameter value, $\phi$, changes over time. A sample every 2.5 hours (2.4 samples per period) is used, which is considered a low sampling regime here. Other simulation parameters: the control input is actuated once an hour; $N = 10$hours, containing ten input steps; $\frac{W}{\Delta t} = 50$; 0.0158 is the expected variance of parameters; $M = 5000$, 0.5 re-sampling threshold. a) The system's output tries to follow the sine wave reference, using the control input outlined in Section 3.2. b) The parameter fitting of $\hat{\phi}$, compared to the true parameters, $\phi$. Each parameter has been normalised by the initial parameter estimation in Section 3.2.

#### S10.4 Varied Sampling Rate Identification And Controls

The identification *MAE*, Eq.(7), indexing the error in the prediction trajectories and the Root Mean Squared Error, control *RMSE* Eq.(9), indexing the controller performance, using the reference described in Section 3.2.2 are again compared over a range of sampling rates in Fig. S12.

For the state trajectories in Fig S12A, there is a transition at a sampling time of 1 hour (6 samples per period), where the identification *MAE* of the kernel filter drops below that of the particle filter, showing that when sampling less than once an hour, the kernel filter outperforms the particle filter. Sampling less than once every 10 hours (0.6 samples per period), both filters tend to have the same identification *MAE*, that of a static model where the parameters are fixed at the initial value. This is because neither filter varies the parameters far from the initial estimation, and the initial steady state oscillation is unaltered by either filter. These features match the three-parameter simulation in Fig. 9A.

The two filters' control Root Mean Squared Error, *RMSE*, is compared over a range of sampling rates in Fig. S12B. There is a transition at 1.25 hours between samples (5.14 samples per period), where the control *RMSE* of the kernel filter drops below that of the particle filter, showing that the kernel filter outperforms the particle filter for sampling times higher than 1.25 hours. This transition point (sampling once an hour) is very similar to the filtering simulation shown in Fig. S12A.

Fig. S12C displays the control *RMSE* over varied sampling rates and prediction horizons, $N$. It can be seen that there is a darker region for the kernel filter compared to the particle filter between 3 and 6 hour sampling times (2-1 samples per period) over all prediction horizons. As in Fig. S12B (for $N = 10$) between sampling times of 6-10 hours, the control *RMSE* of the two filters is similar and, therefore, difficult to distinguish between the colour of the different filters' heatmaps in Fig. S12C.

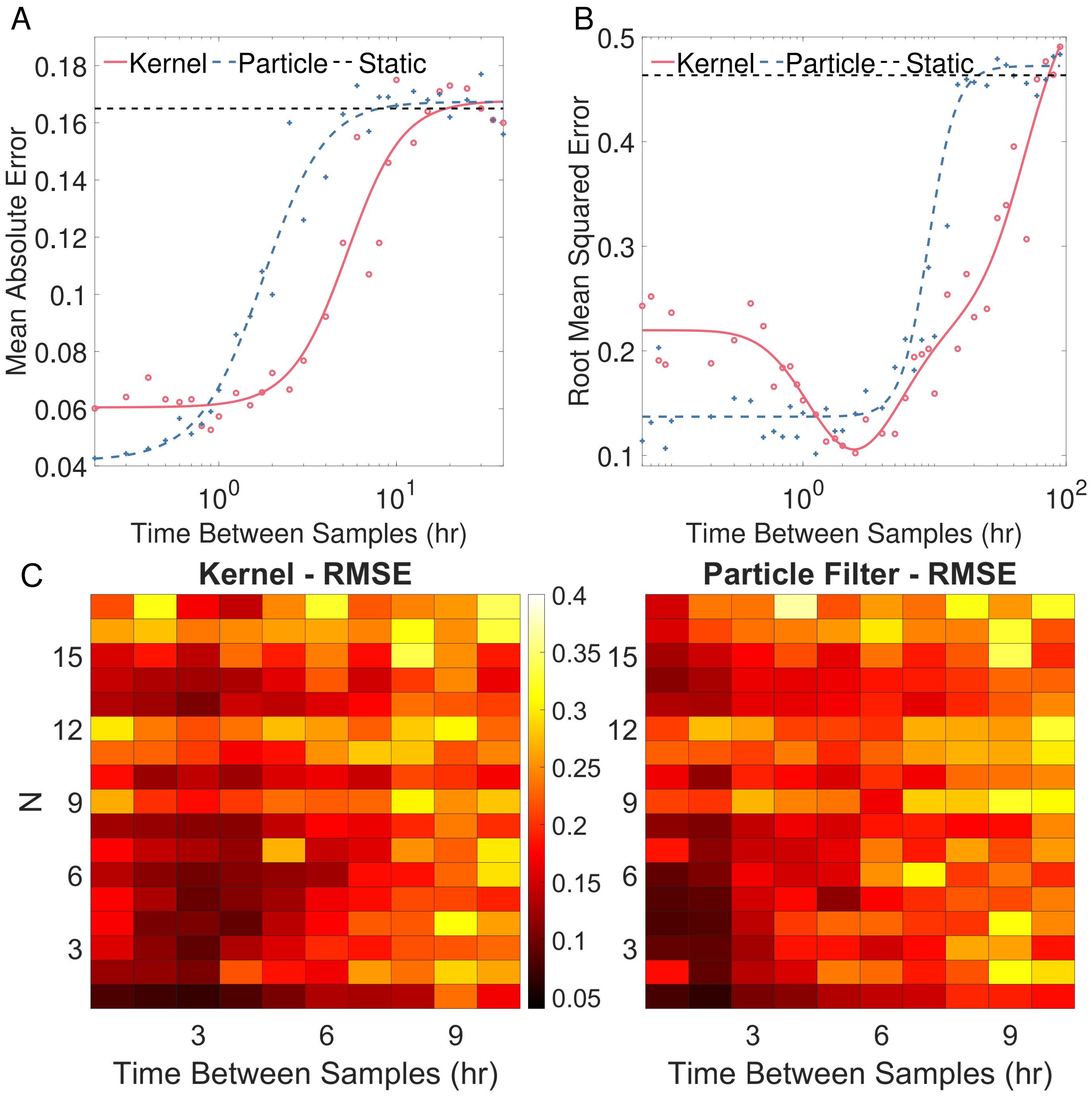


Figure S12: a) The identification Mean Absolute Error (*MAE*, Eq.(7)) of the two filters is compared over a range of sampling rates for the P53 oscillator. The simulation parameters for the kernel and particle filters are noted in Fig S10. b) The control *RMSE* (Eq.(9)) of the two filters are compared over a range of sampling rates for the P53 oscillator, with simulation parameters as in Fig S11. c) The control *RMSE* of the two filters, compared over a range of sampling rates and prediction horizons, *N*, for the P53 oscillator. Other simulation parameters as in Fig S11.

### S11 The Effect Of Added Fitted Parameters On Controller Performance

Several system parameters could have been allowed to vary within the filters. Section 3.2 simulates three mass action constants, and Section S10 looks into varying all six. Fig. S13 displays the control *RMSE* heat maps for varied prediction horizons and sampling times, as described in Figure 9, showing the effect of including more parameters within the filters.

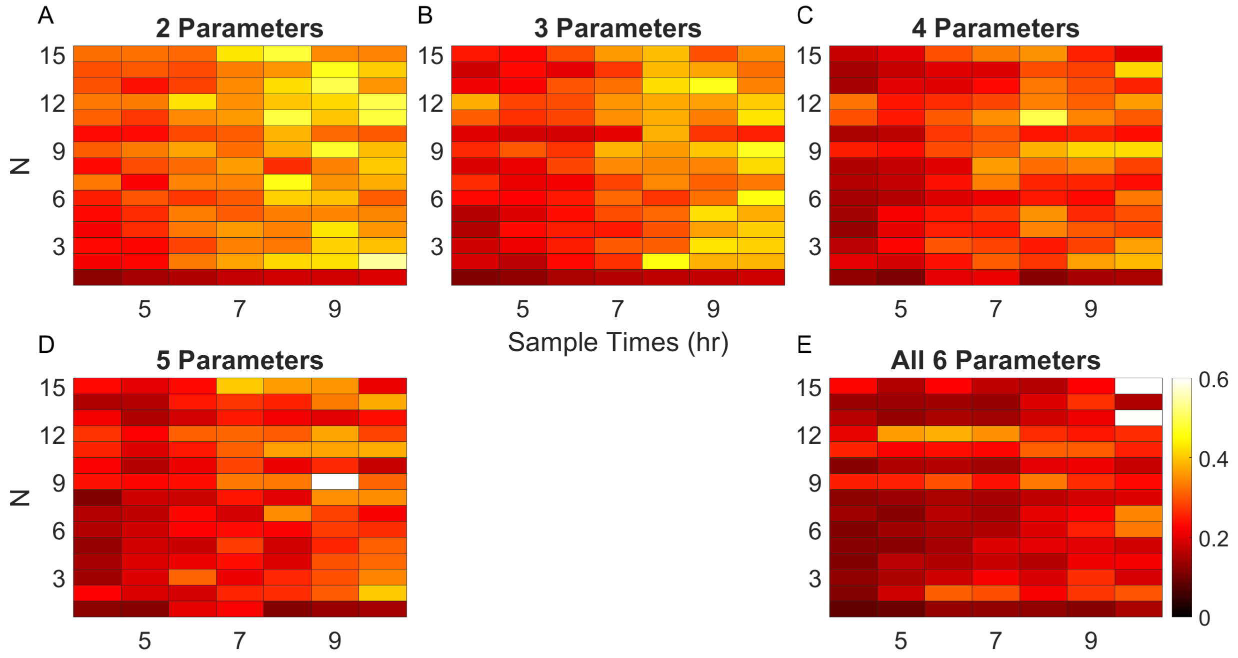


Figure S13: The control *RMSE* heatmaps for varied prediction horizons and sampling times show the effect of including more fitted parameters within the kernel filter. To compare $KC_1 = 0.01$, which is used for the whole window shown. a) Varying the two parameters $\alpha_0$ and $\alpha_k$. b) Varying the three parameters $\alpha_0$, $\alpha_k$ and $\beta_y$ as in Section 3.2. c) Varying the four parameters $\alpha_0$, $\alpha_k$, $\beta_y$ and $\beta_x$. d) Varying the five parameters $\alpha_0$, $\alpha_k$, $\beta_y$, $\beta_x$ and $\alpha_y$ (all parameters except $k$). e) Varying all six parameters, as in Section S10.

In all five simulations in Fig. S13, it can be seen that the bottom left corner is the darkest section of the plot, as discussed in both Sections 3.2.3 and S10.4. It can also be seen that the more parameters included in the filter, on average, the lower the control *RMSE* throughout the plot, as the average control *RMSE* decreases and the plots become darker. However, as the number of parameters increases, the number of anomalous 'light spots' also increases, with one white square with five parameters and two white squares with six parameters.

The larger the number of parameters included in the filter, the filtering problem becomes increasingly ill-conditioned (as discussed in Section S10.2). The kernel filter's grid search is currently limiting its performance. Adding more parameters to the filter increases the number of parameter combinations that provide satisfactory state estimation.

However, this rudimentary nonlinear solver also limits the parameter space that the filter can jump between each time step. Therefore, the kernel filter can be trapped in a local minimum for which performance degrades progressively. Without a more global optimisation algorithm, the filter cannot leave this neighbourhood to retrieve more optimal solutions, leading to the 'light spots'.

In the future, a better nonlinear solver could retain the average reduced control error whilst not leading to the occasional 'light spot'. Three parameters were chosen for Section 3.2 as it had a good trade-off between average control error and control error variance, showing the prediction horizon and sampling time trends without being distorted by 'light spots'.

### S12 The Effect Of The Grid Search On Controller Performance

The grid search outlined in Section 2.1 trials three different values for each fitted parameter, one at the previous optimal parameter value, $\hat{\phi}_{t-1}$, and another two at $\pm\rho\%$. For a system containing $n$ parameters, the discrete set, $\Phi_t$, contains all $3^n$ possible combinations of $\tilde{\phi}_t$ where each element of $\hat{\phi}_{t-1}$ has been multiplied by an element of $([1-\rho, 1, 1+\rho])$ to create a $3^n$ unique trial $\tilde{\phi}_t$. The sample set that produces the highest HSIC will be chosen as the kernel filter's parameter estimate, $\hat{\phi}_t$.

As discussed, this simple grid search is needed over a continuous nonlinear solver so that the MPC can fit its model parameters online and apply the optimal input between each sampling time. However, the limitations of this search have been discussed in Sections 3.2.3 and S11, showing that this discrete approximation for a nonlinear solver leads to a sub-optimal choice in $\hat{\phi}$ and an increase in control error. There is a balance between solver run-time and accuracy.

To show the limiting effect of the current grid search, a 'light spot' of the kernel filter in Fig. 9 ($N = 9$, 2 hours between samples) has been selected to compare the control performances achieved when using two different grid sizes. The 'Small Grid' considered here is the same as the grid considered in Section 2.1 where each element of $\hat{\phi}_{t-1}$ has been multiplied by an element of $([1-\rho, 1, 1+\rho])$ to create a $3^n$ unique trial $\tilde{\phi}_t$. A second, 'Large Grid' comprising $9^n$ sample points and allowing the kernel filter to jump $\pm 4\rho$ from the current parameter values is considered for Comparison, where each element of $\hat{\phi}_{t-1}$ has been multiplied by an element of $([1, 1-\rho, 1+\rho, 1-2\rho, 1+2\rho, 1-3\rho, 1+3\rho, 1-4\rho, 1+4\rho])$, to create $9^n$ unique trials of $\tilde{\phi}_t$. Fig. S14 shows that the control performance is lower for the small grid (control $RMSE = 0.3473$) compared to the larger grid (control $RMSE = 0.2365$), showing that the quality of the optimisation achieved by the grid search method can be a limiting factor. Unfortunately, this increase in performance comes at a cost; the grid now includes nine samples per parameter rather than 3. The solver will take three times longer to find a solution per parameter ($3^n$ grid samples vs $3^{2n}$ grid samples). For the P53 simulation in Fig. S14, the 'Small Grid' simulations took 34s, whereas the 'Large Grid' simulation took 256s. Therefore, the MPC user must find a balance between solver accuracy and run-time when using the kernel filter. The kernel filter can be updated as faster nonlinear solvers are developed, and its control performance will only increase.

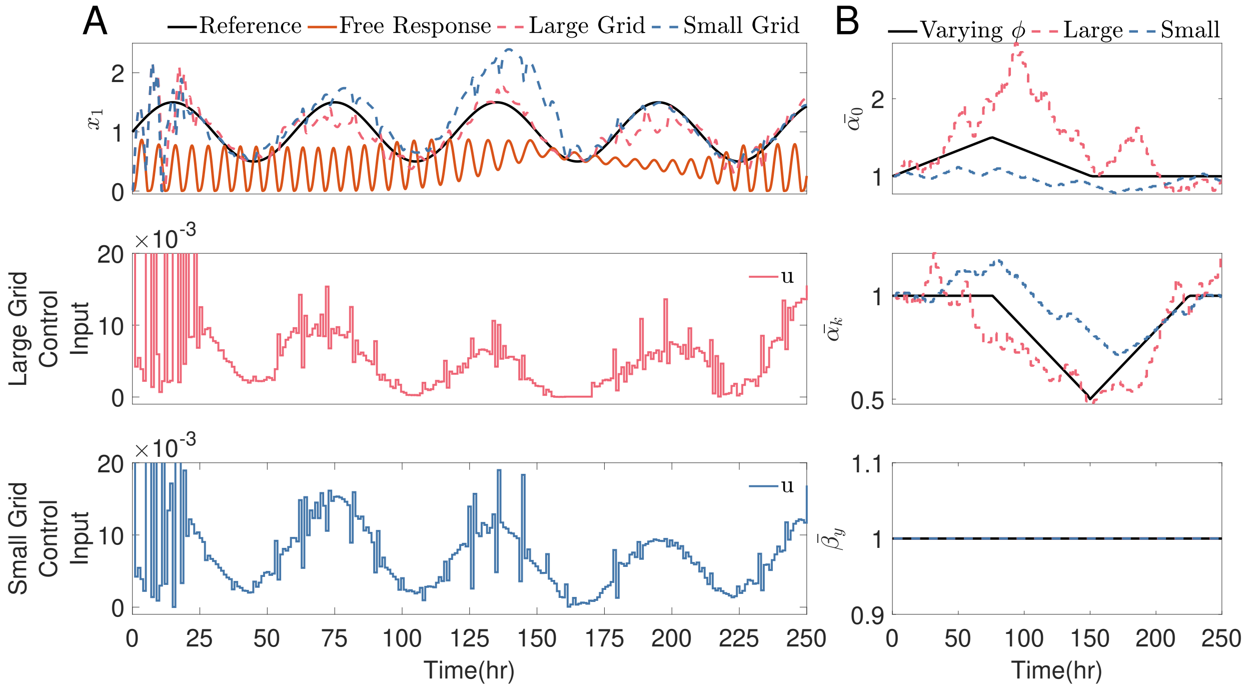


Figure S14: The kernel filter fitting the parameters, online, to match the states of the P53 system (Eq.(13) - Eq.(15)) in which the true parameter value, $\phi$, changes over time. Two size grid searches are compared, as explained in Section S12. A sample every 2 hours (3 samples per period) is used, which is considered a low sampling regime here. Other simulation parameters: the control input is actuated once an hour; $N = 9$ hours, containing nine input steps; $\frac{W}{\Delta t} = 50$. a) The system's output tries to follow the sine wave reference, using the control input outlined in Section 3.2. b) The parameter fitting of $\hat{\phi}$, compared to the true parameters, $\phi$. Each parameter has been normalised by the initial parameter estimation in Section 3.2.

### S13 Toggle Switch

The first four states in the toggle switch model described by Eq. (17) to Eq. 22 are the mass action kinetics of the toggle switch system, containing the TetR and LacI proteins and their respective mRNA molecules. The final two states in the system are new to [9], adding the active transport dynamics of the two inputs into the E. coli cell, differentiating between the extracellular and the intracellular concentration of each input. The inputs are actively transported into/out of the cell; therefore, the rate of transport changes depending on whether there is more of each drug inside or outside the cell, as seen in conditions of states five (Eq. (21)) and six (Eq. (22)).

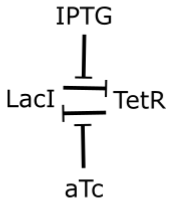


Figure S15: Toggle switch

#### S13.1 Parameters in the Toggle Switch Model

The system has 22 kinematic constants, all described in Table S7. As discussed in Section S11, the filters would be computationally heavy and ill-posed if all 22 of the system parameters were fitted within $\hat{\phi}$. We started from the toggle switch parameter sensitivity and variation analysis between data fitting within the thesis [10]. The parameters $[\eta_{Lac}, \eta_{Tet}, \theta_{Lac}, \theta_{Tet}]$ contain the highest variations between fittings of the ODE models and will form the fitted parameter set $\phi$. These Hill term parameters have been chosen as they characterise the speed of the two cascade reactions within the cell, how TetR inhibits LacI and how LacI inhibits TetR. None of these parameters characterise how the inputs defuse into the cell or how these inputs interact with the two proteins. For all time varying parameter simulations, $\phi_0 = [\eta_{Lac}, \eta_{Tet}, \theta_{Lac}, \theta_{Tet}] = [2.00, 2.00, 31.94, 30.0]$. The state vector is $\mathbf{x} = [\text{mRNA}_{\text{LacI}}, \text{mRNA}_{\text{TetR}}, \text{LacI}, \text{TetR}, \text{aTc}, \text{IPTG}]^T$. The initial conditions $\mathbf{x}_0 = [0.032, 0.119, 50, 1500, 0, 1]^T$ are used in all simulations. The system is used to model experimental data rather than simulations, and therefore, measurement noise is already included in the measured states, and no noise is added to the data.

| Variable | Definition | Value |
|---|---|---|
| $\kappa_{m0}^{L}$ | mRNA baseline transcription rate of LacI's mRNA. | $0.032 min^{-1}$ [9] |
| $\kappa_{m}^{L}$ | mRNA transcription rate of LacI's mRNA. | $8.3 min^{-1}$ [9] |
| $\theta_{a\text{Tc}}$ | Half saturation concentration of *a*Tc. | 31.94 ng.mL$^{-1}$ [9] |
| $\eta_{a\text{Tc}}$ | Hill's coefficient of *a*Tc. | 2.00 [9] |
| $\theta_{\text{TetR}}$ | Half saturation concentration of TetR. | 30.00 a.u. [9] |
| $\eta_{\text{TetR}}$ | Hill's coefficient of TetR. | 2.00 [9] |
| $g_{m}^{L}$ | Degradation rate of LacI's mRNA. | 0.1386 $min^{-1}$ [9] |
| $\kappa_{m0}^{T}$ | mRNA baseline transcription rate of TetR's mRNA. | $0.119 min^{-1}$ [9] |
| $\kappa_{m}^{T}$ | mRNA transcription rate of TetR's mRNA. | $2.06 min^{-1}$ [9] |
| $\theta_{\text{IPTG}}$ | Half saturation concentration of IPTG. | 0.0906 mM[9] |
| $\eta_{\text{IPTG}}$ | Hill's coefficient of IPTG. | 2.00 [9] |
| $\theta_{\text{LacI}}$ | Half saturation concentration of LacI. | 31.94 a.u. [9] |
| $\eta_{\text{LacI}}$ | Hill's coefficient of LacI. | 2.00 [9] |
| $g_{m}^{T}$ | Degradation rate of TetR's mRNA. | 0.1386 $min^{-1}$ [9] |
| $\kappa_{p}^{L}$ | Translation rate of LacI. | 0.9726 a.u. mRNA$^{-1}min^{-1}$ [9] |
| $g_{p}^{L}$ | Degradation rate of LacI. | $0.0165 min^{-1}$ [9] |
| $\kappa_{p}^{T}$ | Translation rate of TetR. | 1.170 a.u. mRNA$^{-1}min^{-1}$ [9] |
| $g_{p}^{T}$ | Degradation rate of TetR. | $0.0165 min^{-1}$ [9] |
| $k_{\text{in}}^{\text{aTc}}$ | *a*Tc exchange rate into the cell. | $0.162 min^{-1}$ [9] |
| $k_{\text{out}}^{\text{aTc}}$ | *a*Tc exchange rate out of the cell. | 0.0200 $min^{-1}$ [9] |
| $k_{\text{in}}^{\text{IPTG}}$ | IPTG exchange rate into the cell. | 0.0275 $min^{-1}$ [9] |
| $k_{\text{out}}^{\text{IPTG}}$ | IPTG exchange rate out of the cell. | 0.111 $min^{-1}$ [9] |

Table S7: Parameters used in the toggle switch model, including the input parameter. a.u. stands for arbitrary fluorescence units.

#### S14 Toggle Switch Experimental Data

The data is taken from [11], using the fourteen files labelled 'DynStim_1' to 'DynStim_14' within the attached GitHub® repository, referred to here as Exp1 to Exp14. All of the data from the 14 experiments is taken from [11], and the average of the cell lines is compared to the open-loop response of the toggle switch model, Eq. (17) to Eq. 22.

As discussed within [11], the active transport of the inputs into the cell (states five (Eq. (21)) and six (Eq. (22)).) is added in an attempt to account for the time delay between the change in the inputs and the observed change in the outputs. Whilst these additional states add a delay, it can be seen more obviously in Fig. S20, Fig. S21, Fig. S22, Fig. S24, Fig. S27 and Fig. S28, that the toggle switch model reacts to the change in input faster than all of the cell profiles. Therefore, there is still a fault in the model structure.

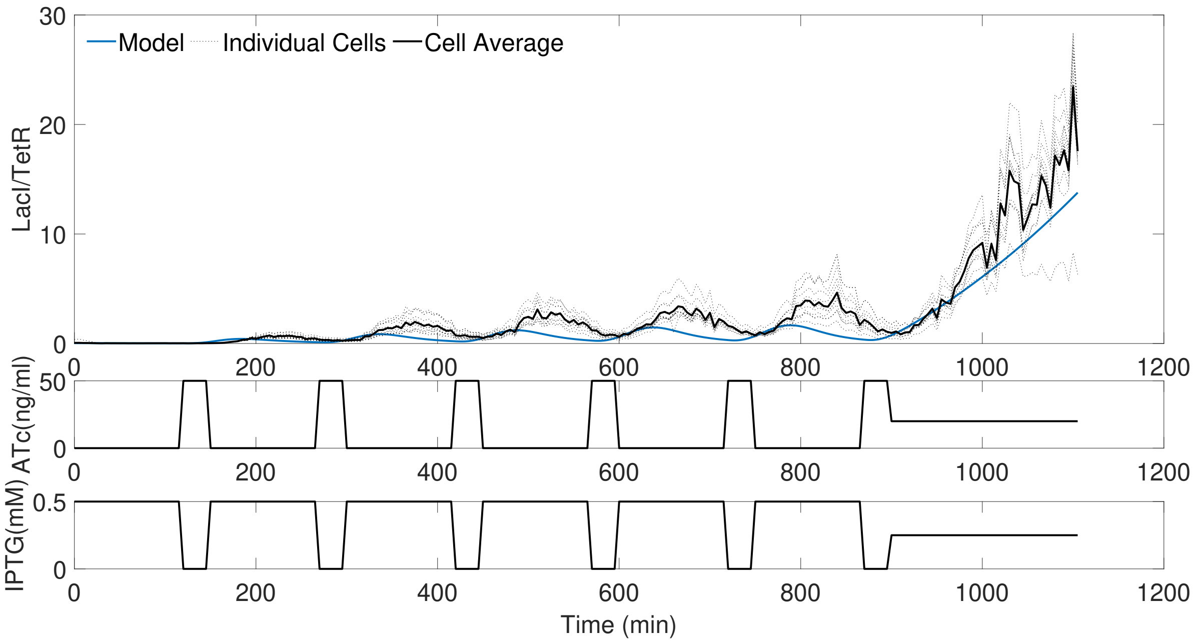


Figure S16: The experimental data within [11], displaying 'DynStim_1', labelled here as Exp1 and comparing the average of the single cells to the toggle switch model in Eq. (17) to Eq. 22 using the parameters from [11], also included in Section S7.

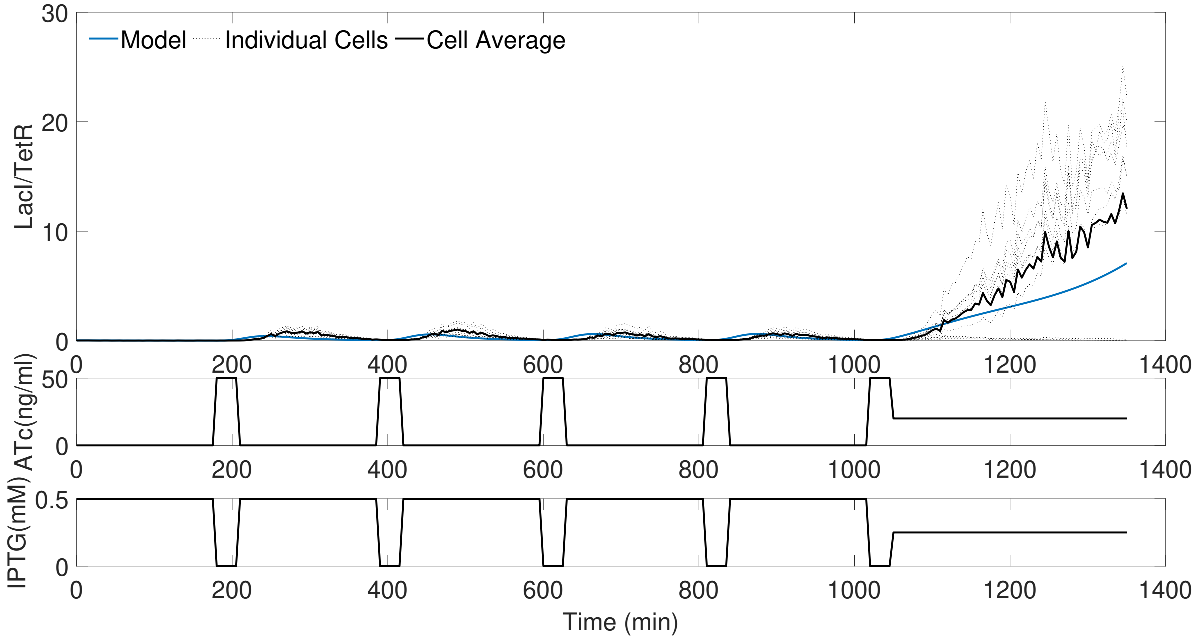


Figure S17: The experimental data within [11], displaying 'DynStim_2', labelled here as Exp2 and comparing the average of the single cells to the toggle switch model in Eq. (17) to Eq. 22 using the parameters from [11], also included in Section S7.

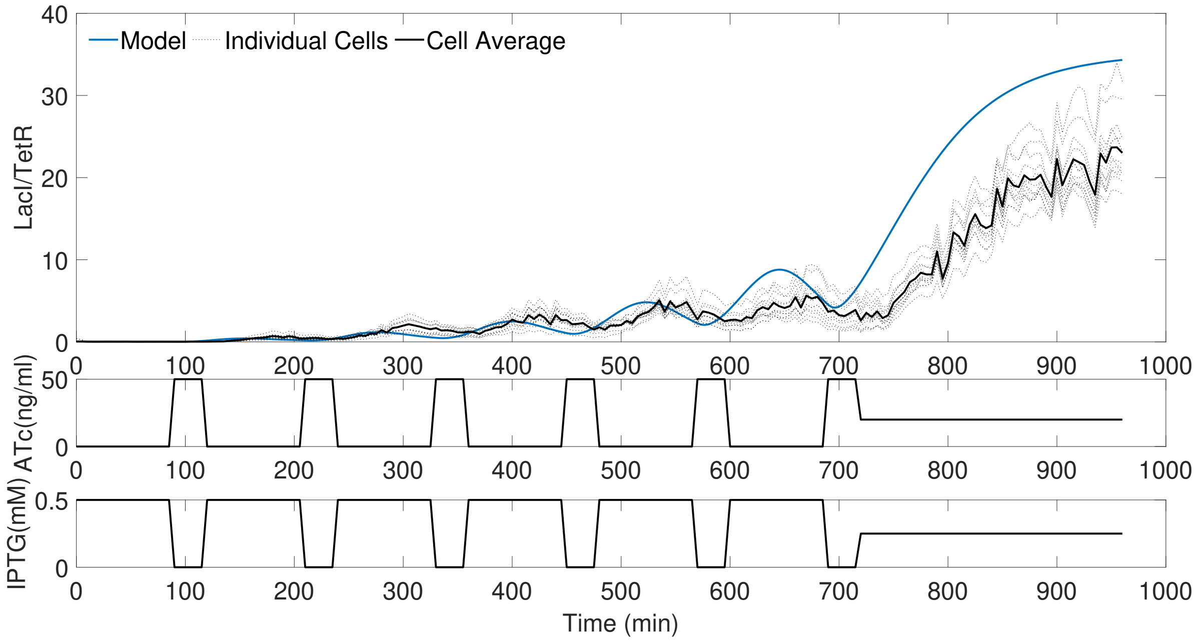


Figure S18: The experimental data within [11], displaying 'DynStim_3', labelled here as Exp3 and comparing the average of the single cells to the toggle switch model in Eq. (17) to Eq. 22 using the parameters from [11], also included in Section S7.

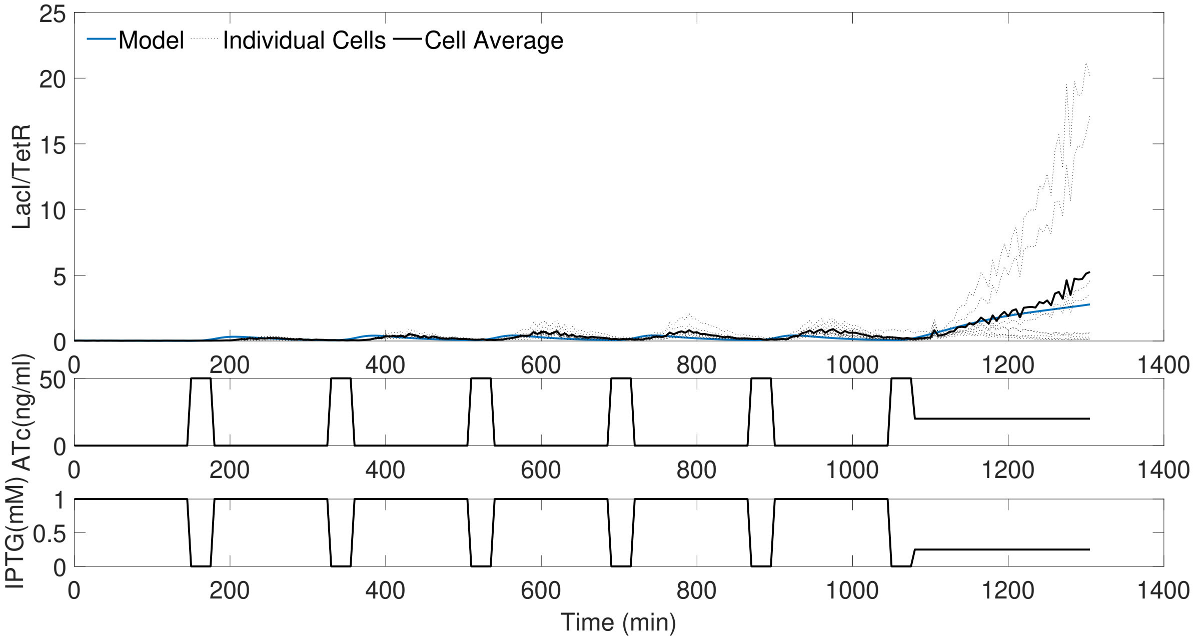


Figure S19: The experimental data within [11], displaying 'DynStim_4', labelled here as Exp4 and comparing the average of the single cells to the toggle switch model in Eq. (17) to Eq. 22 using the parameters from [11], also included in Section S7.

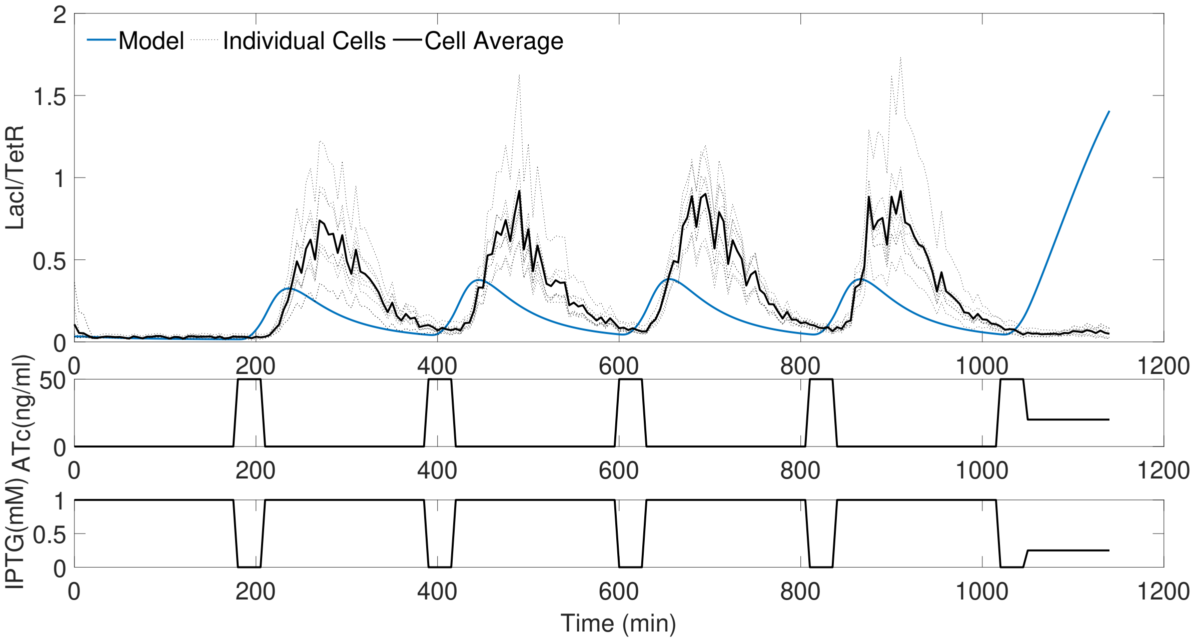


Figure S20: The experimental data within [11], displaying 'DynStim_5', labelled here as Exp5 and comparing the average of the single cells to the toggle switch model in Eq. (17) to Eq. 22 using the parameters from [11], also included in Section S7.

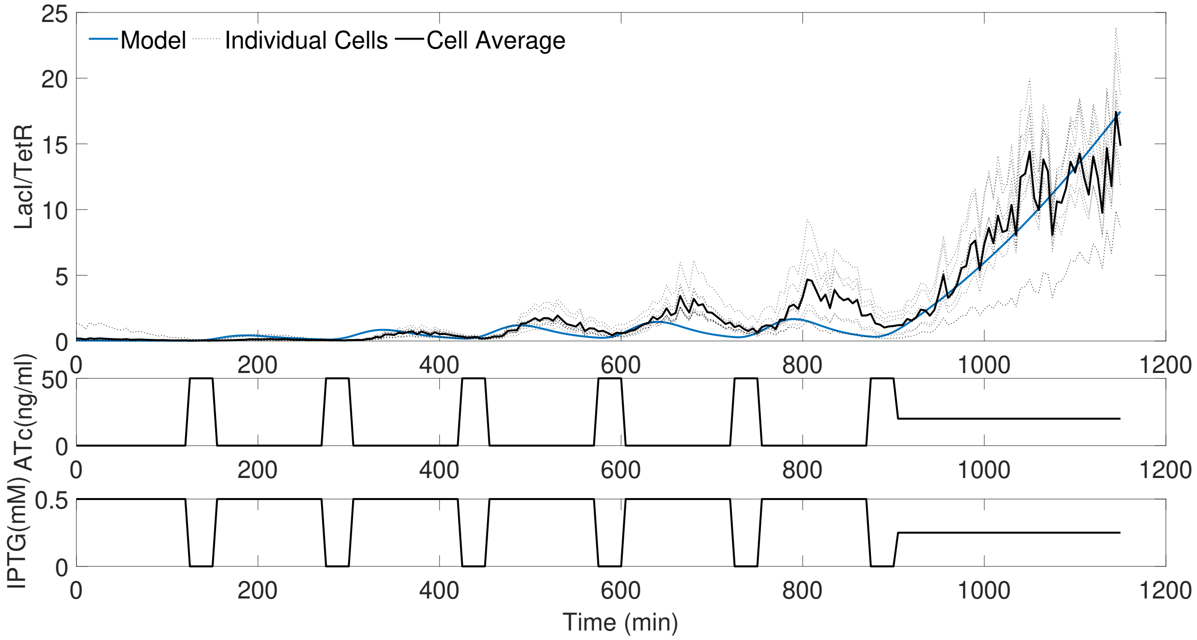


Figure S21: The experimental data within [11], displaying 'DynStim_6', labelled here as Exp6 and comparing the average of the single cells to the toggle switch model in Eq. (17) to Eq. 22 using the parameters from [11], also included in Section S7.

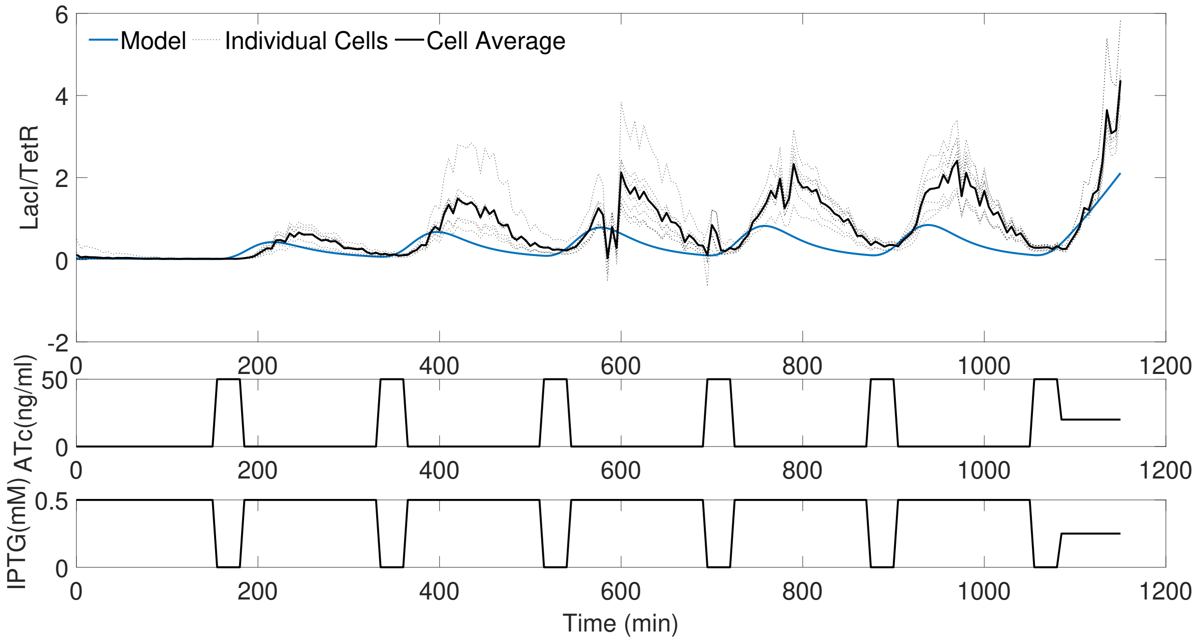


Figure S22: The experimental data within [11], displaying 'DynStim_7', labelled here as Exp7 and comparing the average of the single cells to the toggle switch model in Eq. (17) to Eq. 22 using the parameters from [11], also included in Section S7.

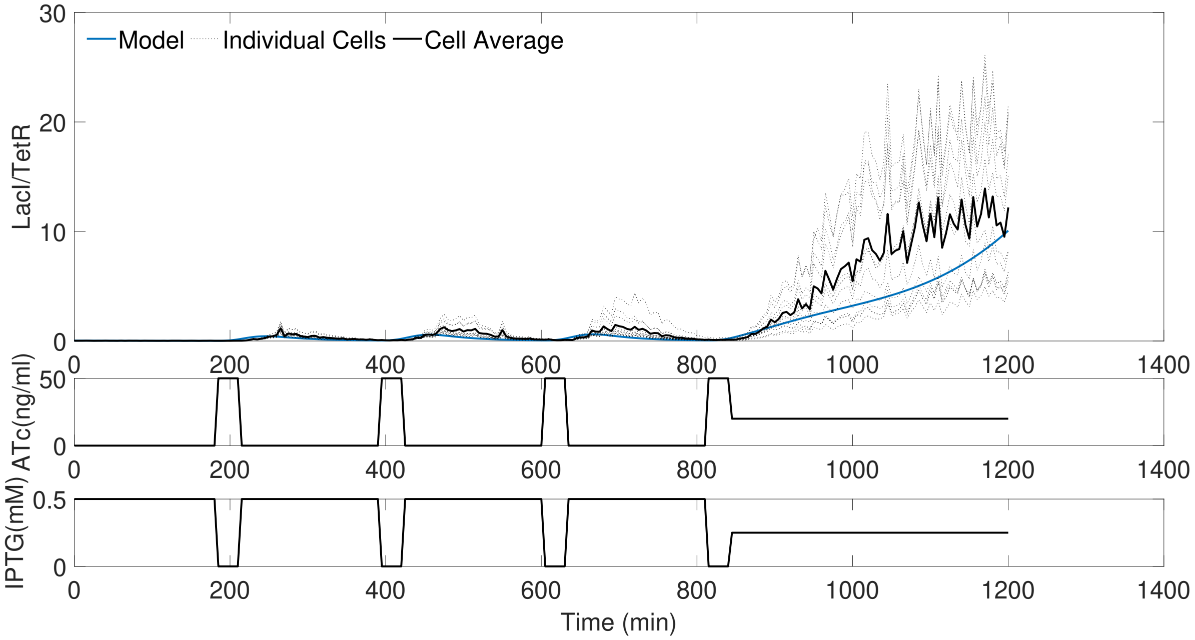


Figure S23: The experimental data within [11], displaying 'DynStim_8', labelled here as Exp8 and comparing the average of the single cells to the toggle switch model in Eq. (17) to Eq. 22 using the parameters from [11], also included in Section S7.

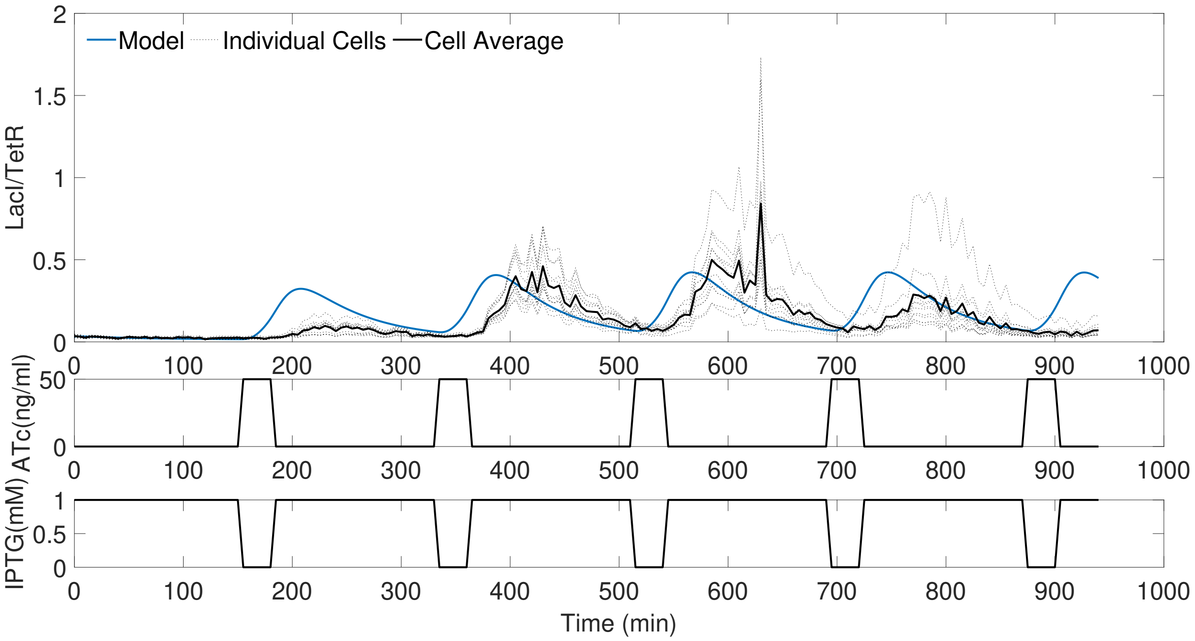


Figure S24: The experimental data within [11], displaying 'DynStim_9', labelled here as Exp9 and comparing the average of the single cells to the toggle switch model in Eq. (17) to Eq. 22 using the parameters from [11], also included in Section S7.

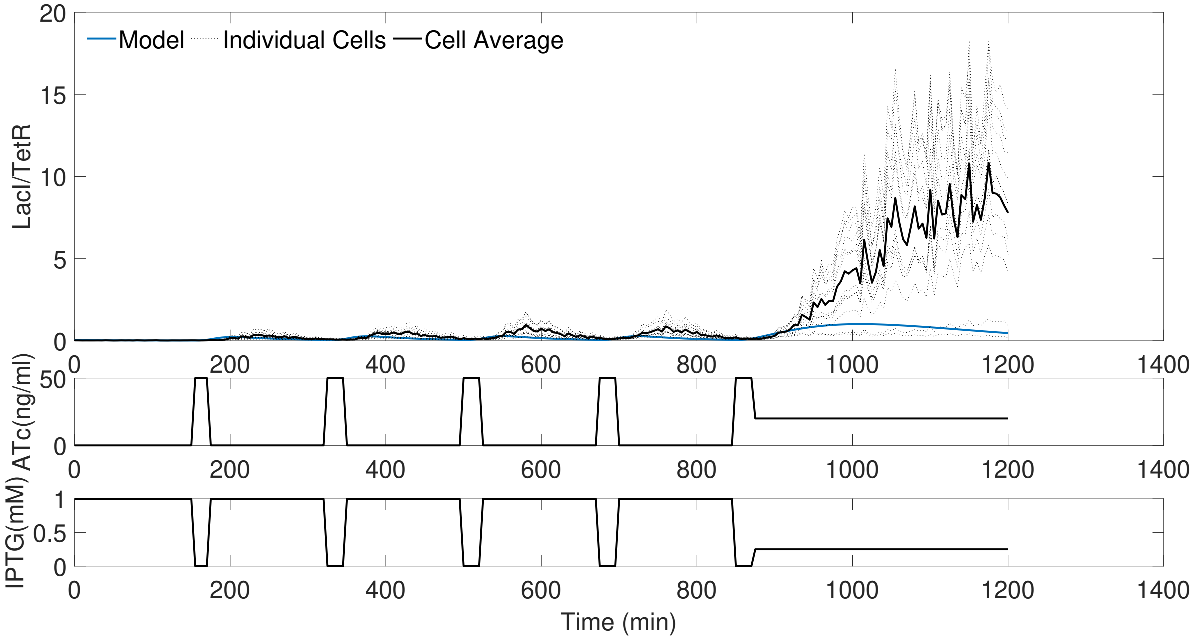


Figure S25: The experimental data within [11], displaying 'DynStim_10', labelled here as Exp10 and comparing the average of the single cells to the toggle switch model in Eq. (17) to Eq. 22 using the parameters from [11], also included in Section S7.

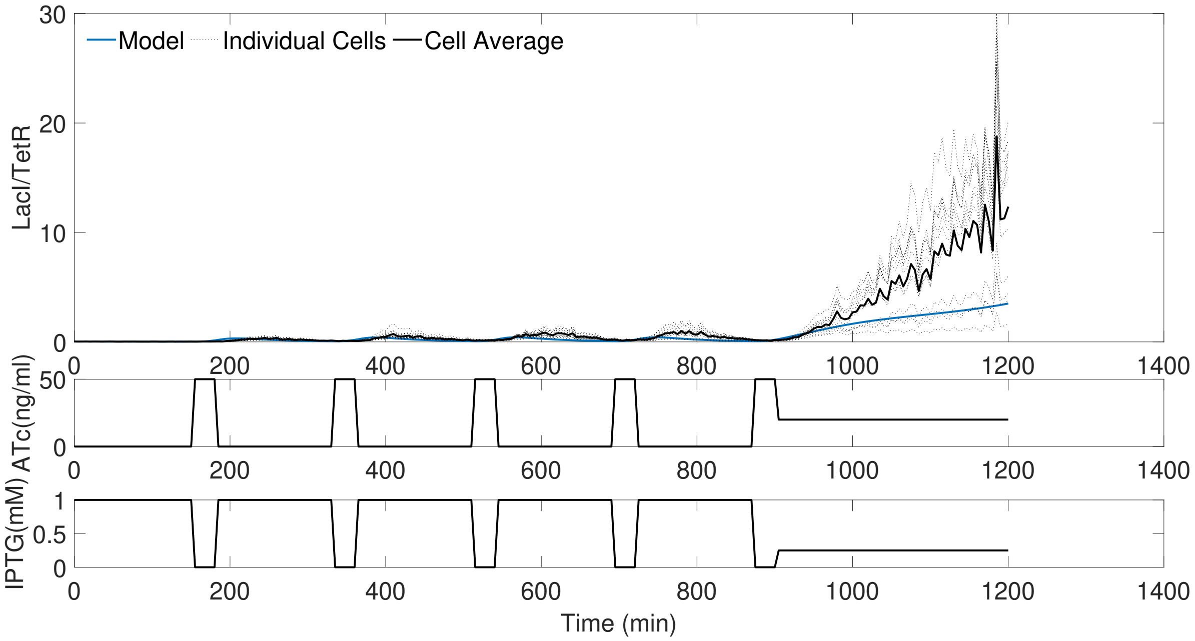


Figure S26: The experimental data within [11], displaying 'DynStim_11', labelled here as Exp11 and comparing the average of the single cells to the toggle switch model in Eq. (17) to Eq. 22 using the parameters from [11], also included in Section S7.

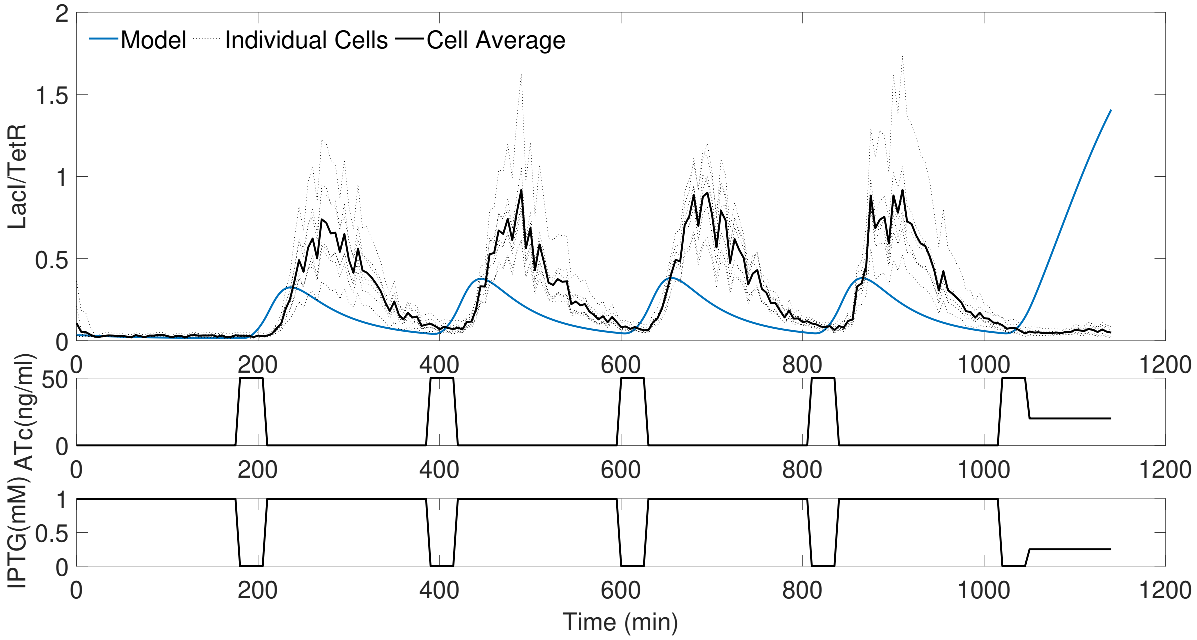


Figure S27: The experimental data within [11], displaying ‘DynStim_12’, labelled here as Exp12 and comparing the average of the single cells to the toggle switch model in Eq. (17) to Eq. 22 using the parameters from [11], also included in Section S7.

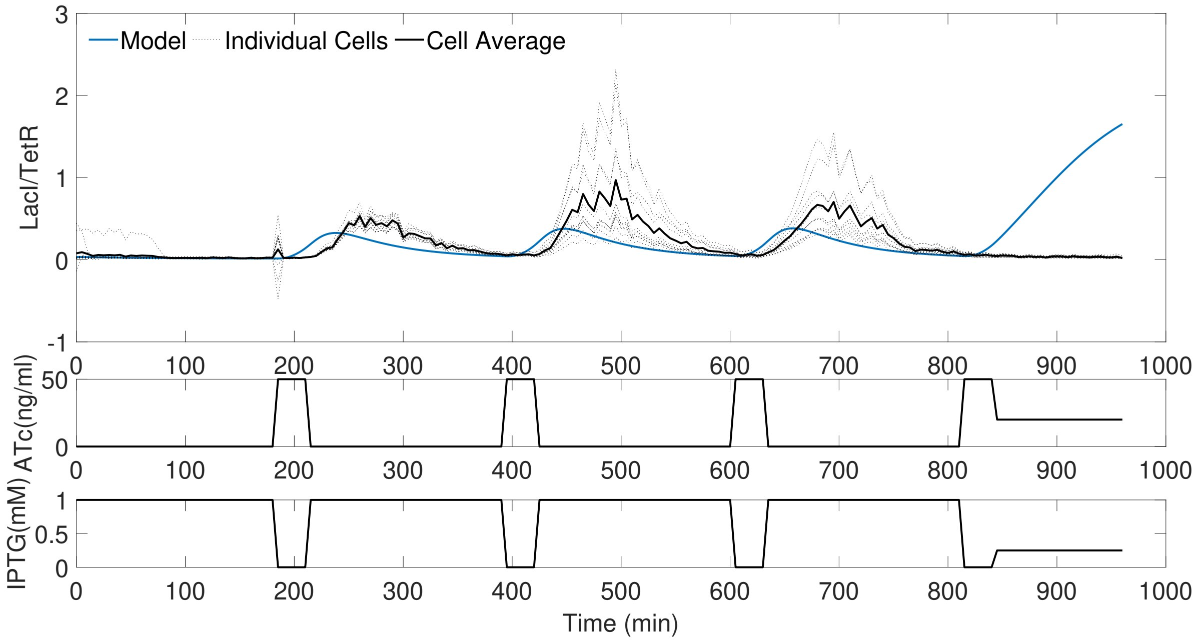


Figure S28: The experimental data within [11], displaying ‘DynStim_13’, labelled here as Exp13 and comparing the average of the single cells to the toggle switch model in Eq. (17) to Eq. 22 using the parameters from [11], also included in Section S7.

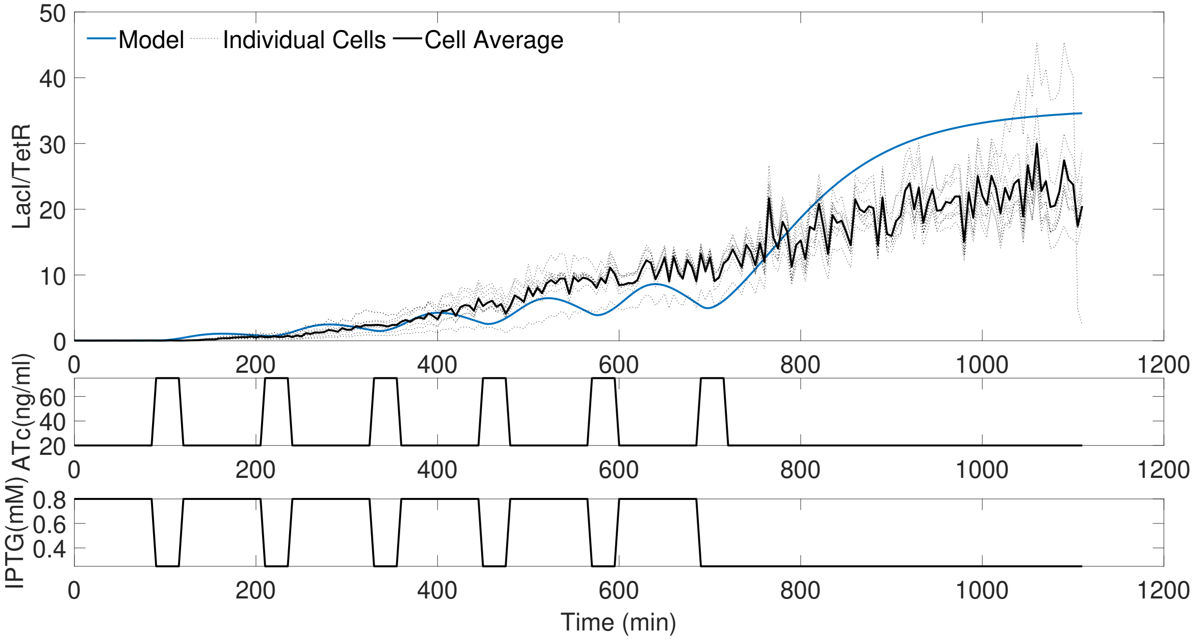


Figure S29: The experimental data within [11], displaying 'DynStim_14', labelled here as Exp14 and comparing the average of the single cells to the toggle switch model in Eq. (17) to Eq. 22 using the parameters from [11], also included in Section S7.

#### S15 Varied Sampling Rate Identification for Each Toggle Switch Experiment

For the synthetic gene oscillator and P53 oscillator, Sections 3.1 and 3.2, several sample rate-dependent tuning guidelines are discussed for the two filters. To show the general performance of the kernel filter and particle filter over a range of different experiments, the filter initial hyperparameters are kept constant, as discussed in Section S3 and Section S4. The identification *MAE* for all 14 dynamic experiments within [11] can be seen in Fig. S30, alongside the average response, also shown in Fig. 12.

It can be seen that the kernel filter outperforms the particle filter on nine out of the 14 experiments within the low sampling regime (5-25 minutes between samples). The kernel filter uses a periodic kernel to relate past measurements to predicted future measurements. This has been shown to work well on periodic systems. However, it can be seen in Fig. S16, Fig. S18, Fig. S21 and Fig. S29, for Exp1, Exp3, Exp6 and Exp14, respectively, that the open-loop response of the toggle switch is not periodic and grows over time. In these four experiments, the static identification *MAE* is significantly greater than the rest of the experiments, as the initial model is periodic. The kernel filter expects a periodic response due to its periodic kernel and does not perform as well as the particle filter in most of these non-periodic experiments (Exp1, Exp3 and Exp14). The particle filter has no memory of past data and only expects periodic data due to the model structure; therefore, on average, it copes better when filtering non-periodic responses. In an experiment with a known non-periodic system, a different kernel family can be selected for the kernel filter that better represents the expected response of the system.

Out of the ten experiments exhibiting periodic responses, the kernel filter outperforms the particle filter eight times within the low sample rate regime, with a higher identification *MAE* in only Exp9 and Exp13. As discussed in Section S3, the kernel hyperparameter that represents the expected time period of the response is set to two hours. The kernel hyperparameter optimisation can tune this value within the first kernel window. Exp9 and Exp13 have relatively large time periods (3 and 3.5 hours, respectively) and contain steps of 1*mM* of IPTG (as opposed to 0.5mM, like some of the other experiments), resulting in small amplitudes of oscillation with large periods. These time periods require the kernel filter's hyperparameter optimisation to significantly change the expected time period, whilst the small amplitude makes it more difficult for the filters to differentiate the experimental noise from the toggle switch outputs.

Therefore, on average, the kernel filter outperforms the particle filter across all of the periodic toggle switch experiments, as can be seen in the average experiment subplot of Fig. S30 and in Fig. 12. If the sample time and time period of the experiment are known in advance, then the initial estimation of the kernel's hyperparameters can be adjusted accordingly, to further improve the kernel filter's identification *MAE*.

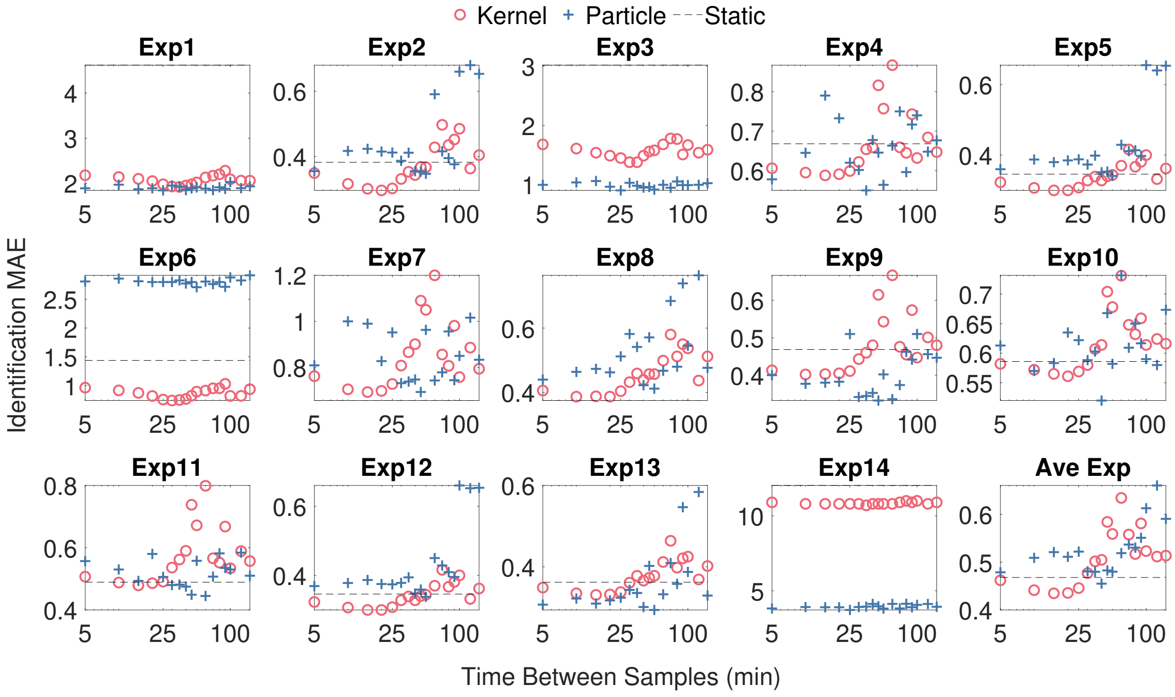


Figure S30: The identification *MAE*, Eq.(7), of the two filters is compared over a range of sampling rates for the toggle switch. The subplots are the identification *MAE* of the 14 experiments (Exp1-Exp14 from [11]) at each time between samples, as labelled. Each experiment used a sample time of 5 minutes (the point on the y-axis). The final subplot is the average identification *MAE* of the periodic experiments (except Exp1, Exp3, Exp6, Exp14). The simulation parameters for the kernel and particle filters are noted in Fig. 11.

[2] Carl Edward Rasmussen and Christopher K. I. Williams. Gaussian Processes for Machine Learning. *Gaussian Processes for Machine Learning*, 12 2018.

[3] David Kristjanson Duvenaud and Pembroke College. Automatic Model Construction with Gaussian Processes Declaration. 2014.

[10] I. De Cesare. *Tools and applications of cybergenetics control of living cells: taming complexity*. PhD thesis, University of Bristol, Bristol, 10 2023.

[11] J. Lugagne et al. Balancing a genetic toggle switch by real-time feedback control and periodic forcing. *Nature Communications*, 2017.
